# Disagreement between demultiplexing methods reveals structured cell quality gradients in multiplexed single-cell data

**DOI:** 10.64898/2026.05.10.724135

**Authors:** Ezgi Sen, Simon Steiger, Mislav Basic, Nina Prokoph, Afzal P. Syed, Isabelle Seufert, Usama-Ur Rehman, Sabrina Schumacher, Anja Baumann, Michaela Feuring, Niels Weinhold, Michael Lübbert, Hartmut Döhner, Konstanze Döhner, Marc S. Raab, Jan-Philipp Mallm, Oliver Stegle, Karsten Rippe

## Abstract

**Background:** Single-cell multi-omics profiling of hematopoietic malignancies frequently involves pooling of patient samples before library preparation to reduce costs. Demultiplexing and quality control of the resulting sequencing data depend on experimental design, sequencing depth, and computational methods. Existing approaches benchmark individual tools, auto-select a single best method, or apply majority voting. However, none systematically exploit disagreement patterns among orthogonal strategies as a diagnostic signal for cell quality.

**Results:** We introduce Split-flow, a modular Nextflow pipeline that runs hashing-based and SNP-based demultiplexing, and transcriptome-based doublet detection in parallel. It classifies cells into quality strata through a concordance-based decision framework. Validation on multiplexed CITE-seq data from 14 multiple myeloma patients across eight Chromium channels demonstrates high reproducibility and shows that discordant cells cluster within specific cell types and quality strata. TCR clonotype cross-referencing against VDJdb confirms that concordance-based classification enriches for biologically genuine immune receptor sequences, with a 5.3-fold enrichment of confirmed public TCR sequences in the high-confidence stratum. Downsampling analysis reveals that SNP-based methods are more depth-sensitive than hash-based approaches, supporting the recommendation to combine both strategies. The framework transfers to AML samples across three assay types (snMultiome-seq, scRNA-seq, scATAC-seq), where ATAC-based demultiplexing resolves donor assignment discordance under low hashing efficiency.

**Conclusions:** Split-flow demonstrates that combining of orthogonal preprocessing methods yields structured information about cell quality and offers a concordance-based framework that transforms this disagreement into a diagnostic signal. It introduces a preprocessing approach that can be exploited beyond hematopoietic malignancies in multiplexed single-cell applications.

**Graphical abstract:** 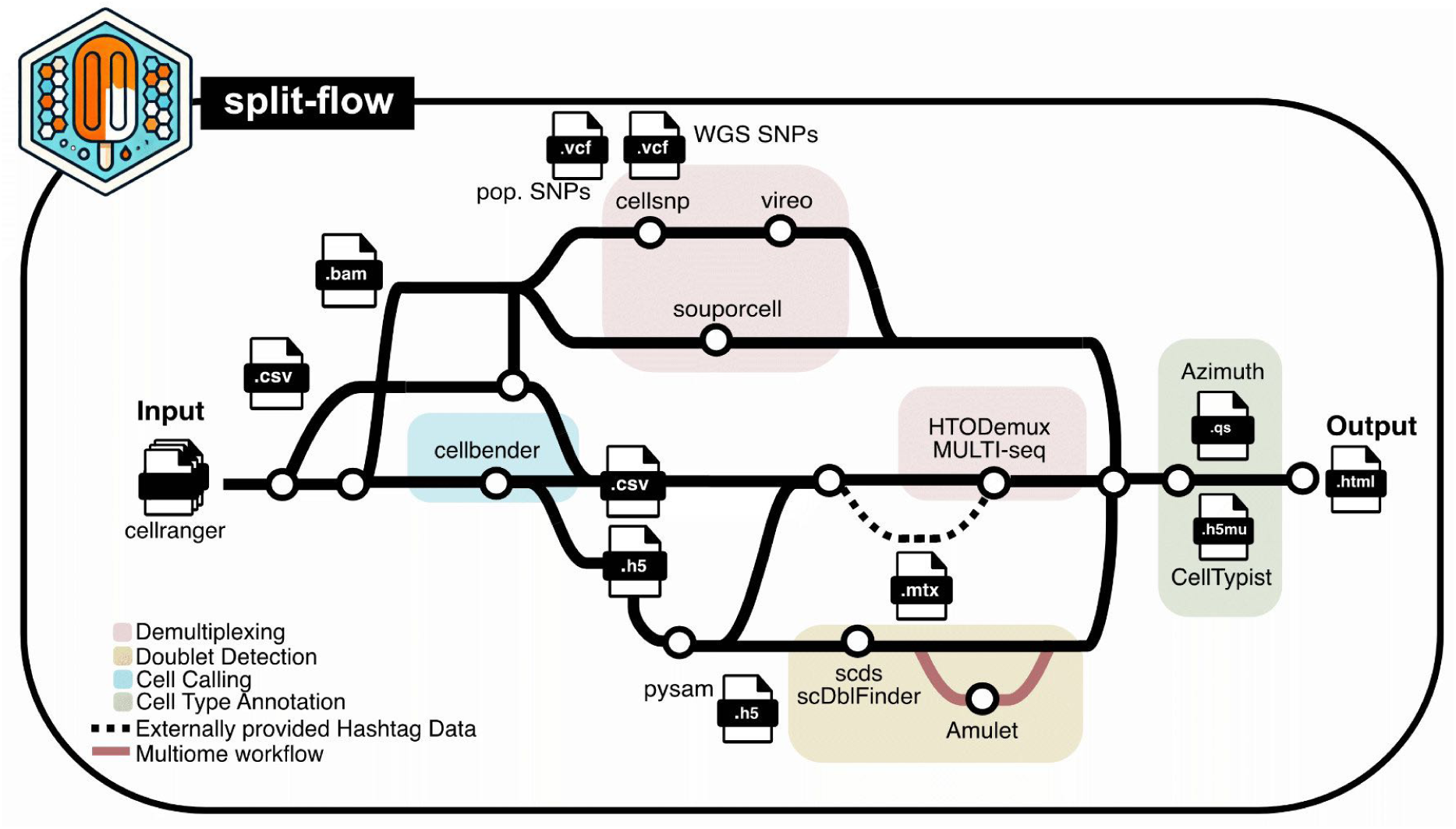

**Highlights and main findings:**

- Introduces Split-flow, a modular Nextflow DSL2 pipeline for preprocessing of multiplexed single-cell multi-omics sequencing data from hematopoietic malignancy samples via a post hoc concordance-based decision framework.
- Provides practical guidance for the experimental design of multiplexed single-cell multi-omics experiments, including the recommendation to combine antibody-based hashing with a SNP genotype reference for orthogonal demultiplexing.
- Reveals that SNP-based demultiplexing is more sensitive to sequencing depth than hash-based approaches, and that the combined strategy mitigates depth-dependent biases in cell-type recovery.
- Demonstrates that disagreement between demultiplexing methods contains structured diagnostic information about cell quality, with concordance categories reflecting genuine quality gradients in multiple myeloma CITE-seq samples.
- Validates the concordance framework using T cell receptor sequences as an orthogonal biological readout, with a 5.3-fold enrichment of confirmed public TCR sequences in the high-confidence stratum.
- Applies the preprocessing framework to AML patient samples across three assay types (snMultiome-seq, scRNA-seq, and scATAC-seq) and demonstrates that ATAC-based demultiplexing can resolve donor-assignment discordance.

## Background

Droplet-based single-cell RNA sequencing (scRNA-seq) enables transcriptome profiling of thousands of individual cells, but the biological insights derived from these data depend critically on how multiplexed samples are preprocessed [1]. Beyond transcriptome profiling alone, droplet-based assays are increasingly combined with additional molecular readouts from the same cells. These include surface protein expression quantified by barcoded antibodies in CITE-seq [2], chromatin accessibility measured by the assay for transposase-accessible chromatin in single cells (scATAC-seq) [3], and immune receptor repertoires resolved by B- and T-cell receptor sequencing (scVDJ-seq) [4]. To jointly assess transcriptional and regulatory information, scRNA-seq has also been integrated with scATAC-seq into combined assays, commonly referred to as scMultiome-seq.

To increase throughput and reduce batch effects, multiplexing strategies were developed for droplet-based platforms [5–9]. However, multiplexing introduces substantial preprocessing complexity, as droplets must be accurately assigned to their sample of origin and multiplets must be identified. These challenges are particularly pronounced in patient-derived samples, which are characterized by variable quality, uneven cell numbers, and limited genotype information. Preprocessing strategies that perform well on simulated data or cell line mixtures often show reduced robustness on real patient material, highlighting the need for workflows that account for experimental design and data limitations.

For multiplexed single-cell sequencing experiments, four conceptually distinct but tightly linked preprocessing steps are cell calling, demultiplexing, doublet detection, and multi-omics integration. These steps are coupled because decisions at each step jointly determine data quality and biological interpretability for downstream analysis.

i. Cell calling distinguishes droplets containing intact cells from empty droplets or debris, using UMI-count thresholds [10–12], ambient RNA modeling [12–14], or joint modeling across modalities as in CellBender [15, 16] or in EmptyDropsMultiome [17]. Different cell calling methods can define different cell sets from the same data, and these initial discrepancies propagate through all subsequent steps.
ii. Demultiplexing assigns cell-containing droplets to their sample of origin, either through hashing-based strategies using sample-specific oligonucleotide labels [9, 18, 19] or through genetic demultiplexing exploiting SNP variation between donors [6]. A growing number of frameworks integrate or compare strategies to improve robustness, including scSNPdemux [20], deMULTIplex2 [21], hadge [22], and YASCP [23]. No single strategy is universally optimal: when multiple methods are applied to the same dataset, they frequently produce different cell-level classifications, and reconciling these discrepancies remains a practical challenge.
iii. Doublet detection identifies droplets containing more than one cell, which can comprise 10–40% of captured droplets depending on loading concentration [24–27]. Current best practice combines demultiplexing-based identification of cross-sample doublets with expression-based methods that infer doublet profiles from transcriptional similarity [24, 25, 28, 29]. However, these orthogonal detection strategies do not always agree on the same cells, and the implications of such discordance for downstream data quality have not been systematically evaluated.
iv. Multi-omics integration must ensure coherent preprocessing of heterogeneous data types from the same cell. Assays such as CITE-seq, scMultiome-seq, or combinations with scVDJ-seq differ in count distributions, sparsity, and noise characteristics [2–4], precluding uniform preprocessing across modalities [15, 16]. When preprocessing methods across modalities yield conflicting quality assessments for the same cell, it remains unclear whether the disagreement should be resolved in favor of one modality or whether the discordance itself carries diagnostic value.

Across all four preprocessing steps, a recurring theme emerges. Multiple computational methods exist for each task. Applying them to the same data frequently produces different cell-level classifications. The prevailing approaches to this cross-method disagreement are to select the best-performing tool, apply majority voting, or filter discordant cells. These strategies share the common assumption that disagreement between methods reflects error to be minimized. Whether the pattern of agreement and disagreement across orthogonal preprocessing strategies carries structured information about cell quality that could improve rather than complicate preprocessing decisions has not been systematically investigated.

Here, we introduce Split-flow, a reproducible Nextflow-based workflow that implements a concordance-based decision framework for multiplexed single-cell sequencing of hematopoietic malignancies. Rather than selecting a single best tool or applying majority voting, Split-flow runs orthogonal preprocessing methods in parallel and uses their agreement and disagreement patterns as diagnostic signals for cell quality. We validate the framework on multiple myeloma (MM) and acute myeloid leukemia (AML) patient samples across CITE-seq, snMultiome-seq, and separate scRNA-seq/scATAC-seq, demonstrating that exploiting cross-method disagreement as a diagnostic signal improves cell selection and reduces systematic bias in downstream biological conclusions.

## Results

### Dual barcoding maximizes preprocessing information for multiplexed single-cell multi-omics of hematopoietic malignancies

A key design question for multiplexed single-cell experiments with clinical samples is how to configure the pooling and barcoding strategy so that preprocessing tools have sufficient orthogonal information to resolve ambiguous cell assignments. We addressed this question using viable frozen bone marrow mononuclear cells (BMMCs) and peripheral blood mononuclear cells (PBMCs) from MM and AML patients, as well as from healthy donors, on the 10x Genomics Chromium platform (**Fig. 1a**).

**Figure 1.**
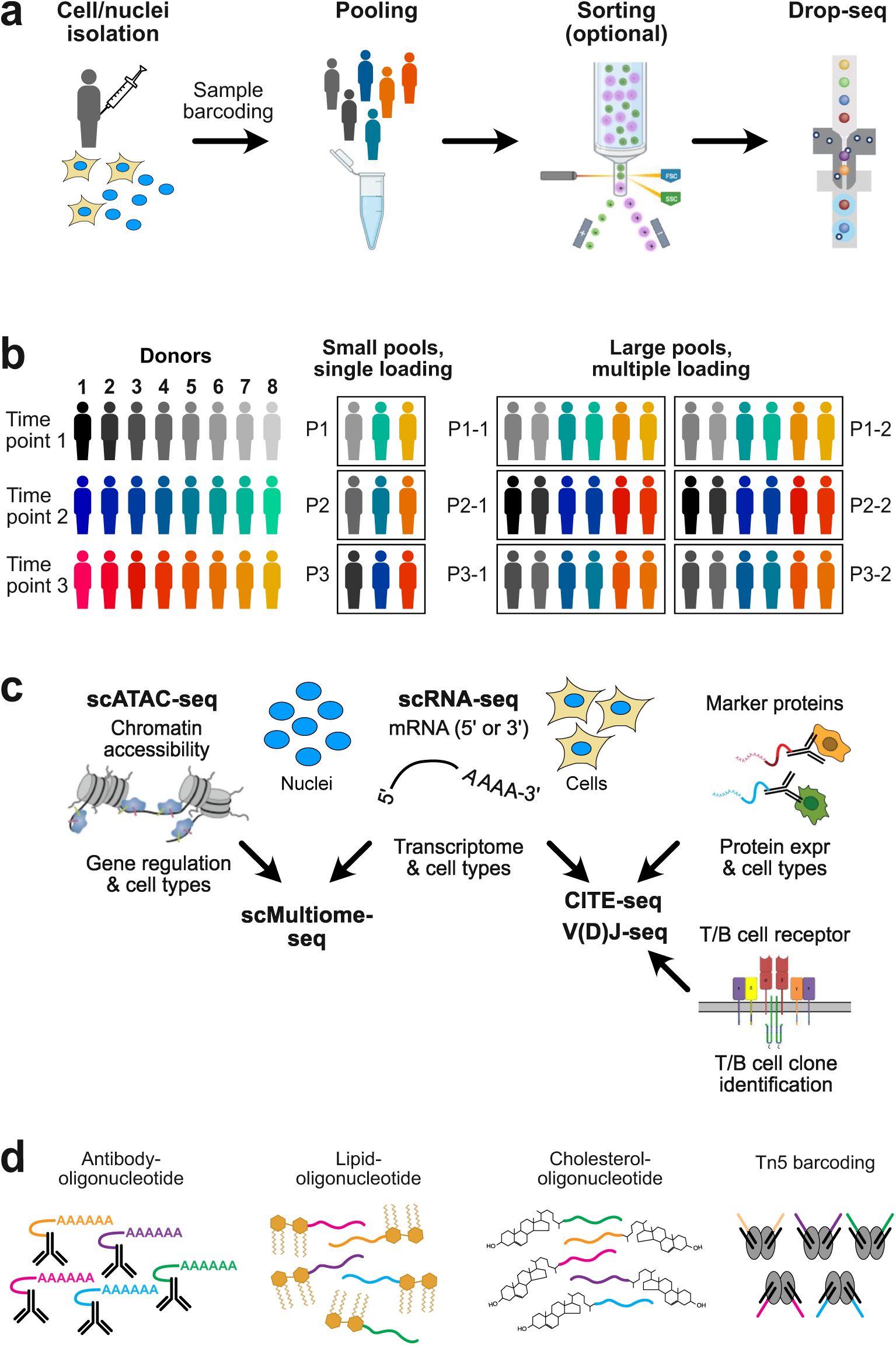
Experimental workflows for multiplexed sc-seq analysis. (**a**) General workflow. (**b**) Hashing techniques. For cell multiplexing, oligonucleotide barcodes conjugated to an antibody or lipid are used [30, 31]. For nuclei, CMOs or barcoding with Tn5 are used. (**c**) Different readouts. (**d**) Pooling of samples.

After thawing, viable cells were enriched by fluorescence-activated cell sorting (FACS). The FACS step can alternatively be performed after sample barcoding and pooling to streamline the workflow and minimize cell/nuclei loss.

The first design choice concerns pooling strategy. Multiplexing enables 3-4 times more cells/nuclei per Chromium channel (∼30-40 k instead of ∼10 k), because cross-donor doublets can be identified computationally. Loading the same pool across multiple channels enables detection of technical variation. Pooling a larger number of genetically distinct samples (e.g., 8-14) and loading multiple times is preferable to pooling only 3-4 samples once (**Fig. 1b**). For the analysis of BMMCs described here, 14 samples were pooled, and ∼38,000 cells were loaded 8 times across the channels of one Chromium flow cell, corresponding to an input of ∼22,000 cells/sample. The second design choice concerns the modality and readout configuration. We optimized preprocessing for three experimental approaches (**Fig. 1c**). For AML samples, we either analyzed snRNA-seq and snATAC-seq separately from the same sample or performed RNA and ATAC analysis simultaneously from the same cell. The CITE-seq analysis was conducted for MM samples using a Total-Seq antibody panel from Biolegend that targeted 192 surface proteins, enabling high-resolution transcriptome analysis and immune cell type profiling. The third and most consequential design choice is the barcoding strategy (**Fig. 1d**). For intact cells, samples can be labeled with antibody-conjugated hashtag oligonucleotides (HTOs) or with lipid-modified oligonucleotides (LMOs) [31, 32]. For nuclei, the validated reagents are cholesterol-modified oligonucleotides (CMOs) [30, 31] and antibodies that target the nuclear pore complex [33]. For snATAC-seq, efficient sample barcoding is achieved by custom Tn5 loading with different sample barcodes, so the sample barcode is introduced during library construction [3]. In this case, sample demultiplexing can be performed directly during library demultiplexing after sequencing.

Hashing-based labeling requires at least ∼100,000 cells for washing steps; for low-input samples, demultiplexing must rely on an external SNP reference (**Table 1**). In our experience, consistent hashing efficiencies above 90% are difficult to achieve, with values of 60-70% not uncommon. SNP-based genetic demultiplexing complements hashing-based approaches by assigning donor identity even when hashing efficiency is low but can fail for cells with low RNA or SNP coverage. We therefore recommend combining hashing or Tn5-based barcoding with SNP-based demultiplexing to maximize cell recovery. An alternative paradigm not evaluated in this study is combinatorial fluidic indexing, exemplified by scifi-RNA-seq [35], which encodes sample and cell identity through plate-based RT preindexing followed by microfluidic droplet barcoding. Critically, the dual-barcoding design recommended here provides two independent lines of evidence for each cell’s donor identity, creating the foundation for a concordance-based preprocessing strategy in which the pattern of agreement and disagreement between methods becomes itself informative.

**Table 1.**
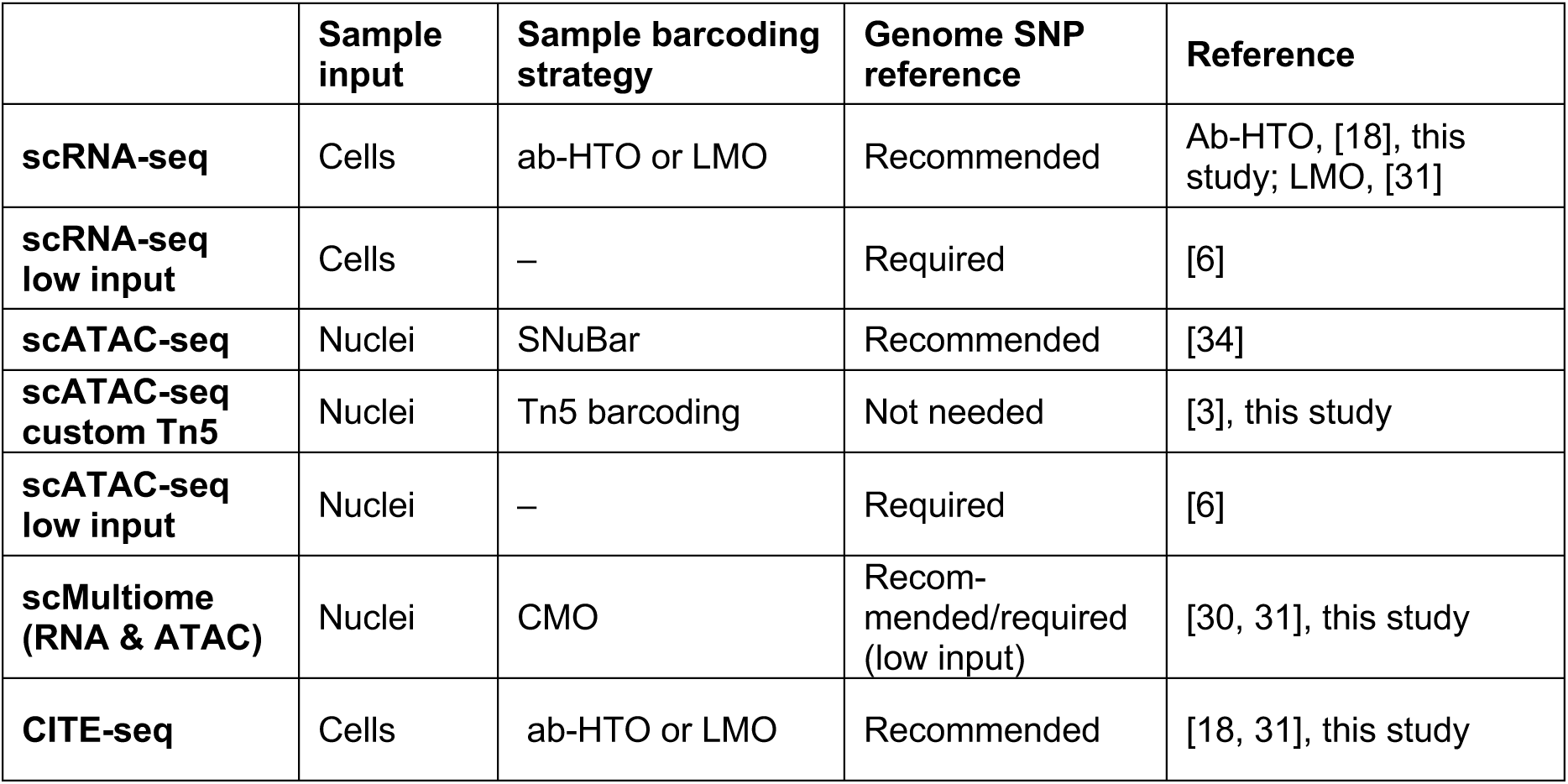
Experimental readouts and sample barcoding strategies.

|  | Sample input | Sample barcoding strategy | Genome SNP reference | Reference |
| --- | --- | --- | --- | --- |
| **scRNA-seq** | Cells | ab-HTO or LMO | Recommended | Ab-HTO, [18], this study; LMO, [31] |
| **scRNA-seq low input** | Cells | – | Required | [6] |
| **scATAC-seq** | Nuclei | SNuBar | Recommended | [34] |
| **scATAC-seq custom Tn5** | Nuclei | Tn5 barcoding | Not needed | [3], this study |
| **scATAC-seq low input** | Nuclei | – | Required | [6] |
| **scMultiome (RNA & ATAC)** | Nuclei | CMO | Recommended/required (low input) | [30, 31], this study |
| **CITE-seq** | Cells | ab-HTO or LMO | Recommended | [18, 31], this study |

### Multiplexed CITE-seq of multiple myeloma samples shows high reproducibility across Chromium channels

To establish that technical variation across Chromium channels does not confound downstream concordance analysis, we first assessed the reproducibility of preprocessing. This strategy was implemented for multiplexed CITE-seq of BMMCs from MM patient samples: 14 patient samples (P1-P14, ∼38,000 cells each) were loaded into eight channels (C1-C8) of a single 10x Chromium flow cell.

UMAP embeddings and recovered cell-type compositions were similar across the eight channels loaded with the same biological samples (**Fig. 2a-b**). Consistent with this, the number of recovered cell barcodes after Cell Ranger processing showed small technical variation (**Fig. 2c**). Cell-containing droplet recovery ranged from 17,083 to 22,182 cells across the eight channels (average 18,954 ± 1,493 cells, CV = 7.9%), with the highest recovery in the last loaded channel. The distribution of cell-resolved QC metrics, including total UMI and ADT counts, unique detected genes per cell, and mitochondrial RNA percentages, was consistent across channels (**Fig. 2c)**. Overall, these results show that data quality was maintained across independent GEM generations (**Supplementary Fig. S2a-d**).

**Figure 2.**
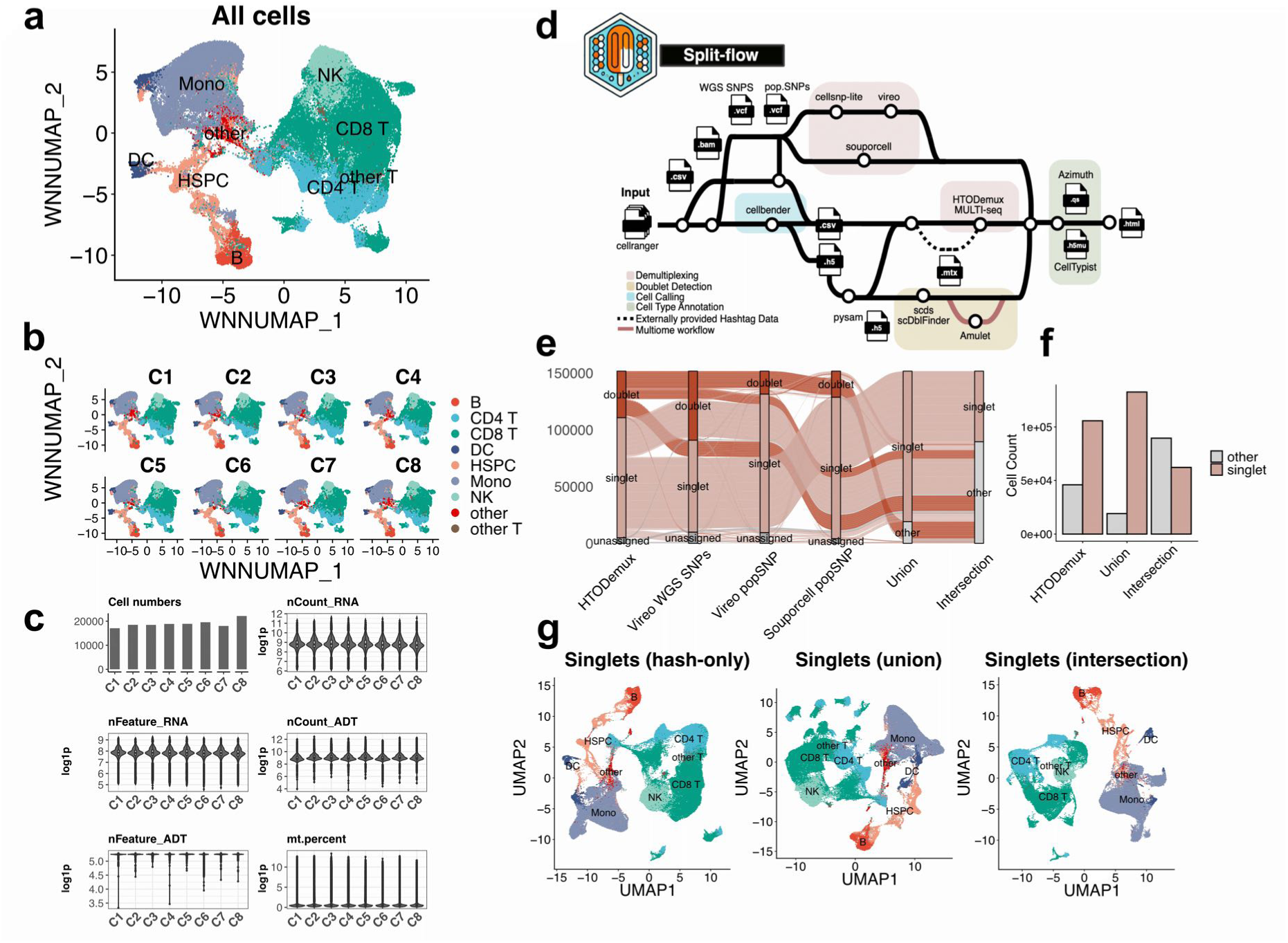
Technical reproducibility, Split-flow pipeline, and methods discordance. **(a)** UMAP embedding based on a weighted nearest neighbor approach that integrates single-cell RNA (transcriptomes) and ADT (surface protein) data for the combined dataset of all CellRanger multi-called cells. **(b)** The same embedding separated by Chromium channels. **(c)** Distribution of QC metrics for all cell-containing droplets across Chromium channels: estimated number of droplets containing cells, total UMI and ADT counts detected, number of unique features for both RNA and ADT, and mitochondrial RNA percentage per cell. Violin plots with overlaid boxplots show the median and interquartile range. **(d)** Overview of the Split-flow preprocessing pipeline. For each Chromium channel, the pipeline sequentially performs multi-modal cell calling and ambient RNA correction with CellBender,demultiplexing using hashes (HTODemux, MultiSeqDemux) and exploiting genetic variation between multiplexed donors (Souporcell, Vireo), transcriptome-based doublet detection (scds, scDblFinder), and cell type annotation either by reference mapping to bone marrow datasets (Azimuth) or logistic regression-based classification (celltypist). Processed data are exported as multimodal objects compatible with both R/Seurat and Python/Scanpy frameworks, and an automated HTML report on the dataset is provided to the user. **(e)** Alluvial plot comparing droplet assignments across HTODemux, Vireo with WGS-derived SNPs, Vireo with population-level SNPs, and Souporcell with population-level SNPs of the multiplexed CITE-seq MM dataset. Additionally, summary categories are added: Union of singlets, defined as droplets that are assigned as singlets by at least one method, and Intersection of singlets, representing droplets classified as singlets by all strategies. All remaining droplets are grouped as others in the summary categories. Flow widths correspond to cell counts; colors indicate HTODemux classifications (singlet, light pink; doublet, dark pink; unassigned, dark grey; others, light grey). **(f)** Bar plots summarizing the absolute counts of singlets and non-singlets identified by hash-based demultiplexing, the union of singlets, and the intersection of singlets. (**g**) Cell-type composition of singlets highlighted on UMAP embeddings computed from SCT-normalized RNA counts for three different sets defined by different demultiplexing strategies (hash-based singlets, union-based singlets, and intersection-based singlets plotted from left to right), annotated using Azimuth with the bone marrow reference (level 1 hierarchy).

### Split-flow integrates parallel demultiplexing with concordance-based quality assessment

We developed Split-flow, a modular Nextflow DSL2 preprocessing pipeline for multiplexed single-cell sequencing data (**Fig. 2d**). For each channel, Split-flow runs CellBender for ambient RNA removal, then performs demultiplexing using hashing-based (HTODemux, MultiSeqDemux) and SNP-based approaches (Souporcell, Vireo) with WGS-derived or population-level reference SNP panels (e.g., [36]), followed by transcriptome-based doublet detection with scds and scDblFinder. Results are exported as multimodal objects in R/Seurat and Python/muon formats. This modular architecture enables systematic comparison of demultiplexing outcomes across methods and provides the foundation for the concordance-based classification described below.

### Hash-based and SNP-based demultiplexing yield substantially different singlet sets

We compared cell-level classifications produced by HTODemux, Vireo (with WGS-derived or population-level SNPs), and Souporcell on the MM CITE-seq data. The alluvial plot in **Fig. 2e** shows that these methods assign different singlet, doublet, and unassigned labels to the same barcodes. To quantify these discrepancies, we defined two summary categories: the union of singlets (labeled as singlets by at least one of the methods) and the intersection (labeled as singlets by all methods) (**Fig. 2e**). Restricting the analysis to the consensus singlet set reduced the number of singlets by 41% (to 62,110) as compared with singlets identified by hash-based demultiplexing alone (105,712 singlets). In contrast, the union-of-singlets strategy increased the total number by 25% (to 132,509) (**Fig. 2f**). Beyond differences in total singlet recovery, the choice of demultiplexing strategy also affected the transcriptional structure captured in the data. When UMAP embeddings were recalculated separately for each singlet set, the union singlet set revealed more detailed structure, especially within T cell populations, where smaller distinct subpopulations became separated in the embedding that were not visible in the other singlet sets (**Fig. 2g**). Overall, these results show that the choice of demultiplexing strategy significantly affects both the number of singlets retained and the transcriptional structures captured in the data. A critical question arising from these observations is whether the discrepancies between methods are randomly distributed across cells or whether they contain structured information about cell identity and quality.

### SNP-based demultiplexing recovers additional singlets but is more sensitive to sequencing depth

To quantify agreement between demultiplexing and doublet-detection methods, we created UpSet plots showing singlet annotations across six methods applied to the multiplexed CITE-seq dataset. In the full-depth dataset, 37.4% of cells were classified as singlets by all four demultiplexing methods and both doublet-detection methods, with concordant donor assignments across all strategies. This percentage increased to 56.5% when the strict requirement for Vireo WGS agreement was relaxed. SNP-based methods consistently labeled a larger proportion of cells as singlets than HTO-based demultiplexing alone (**Fig. 3a**). Among singlets recovered by majority vote, donor assignments were consistent, with only a negligible fraction of cross-donor misassignments. Having established that SNP-based methods recover more singlets at full sequencing depth, we next asked how sensitive this advantage is to reduced coverage.

**Figure 3.**
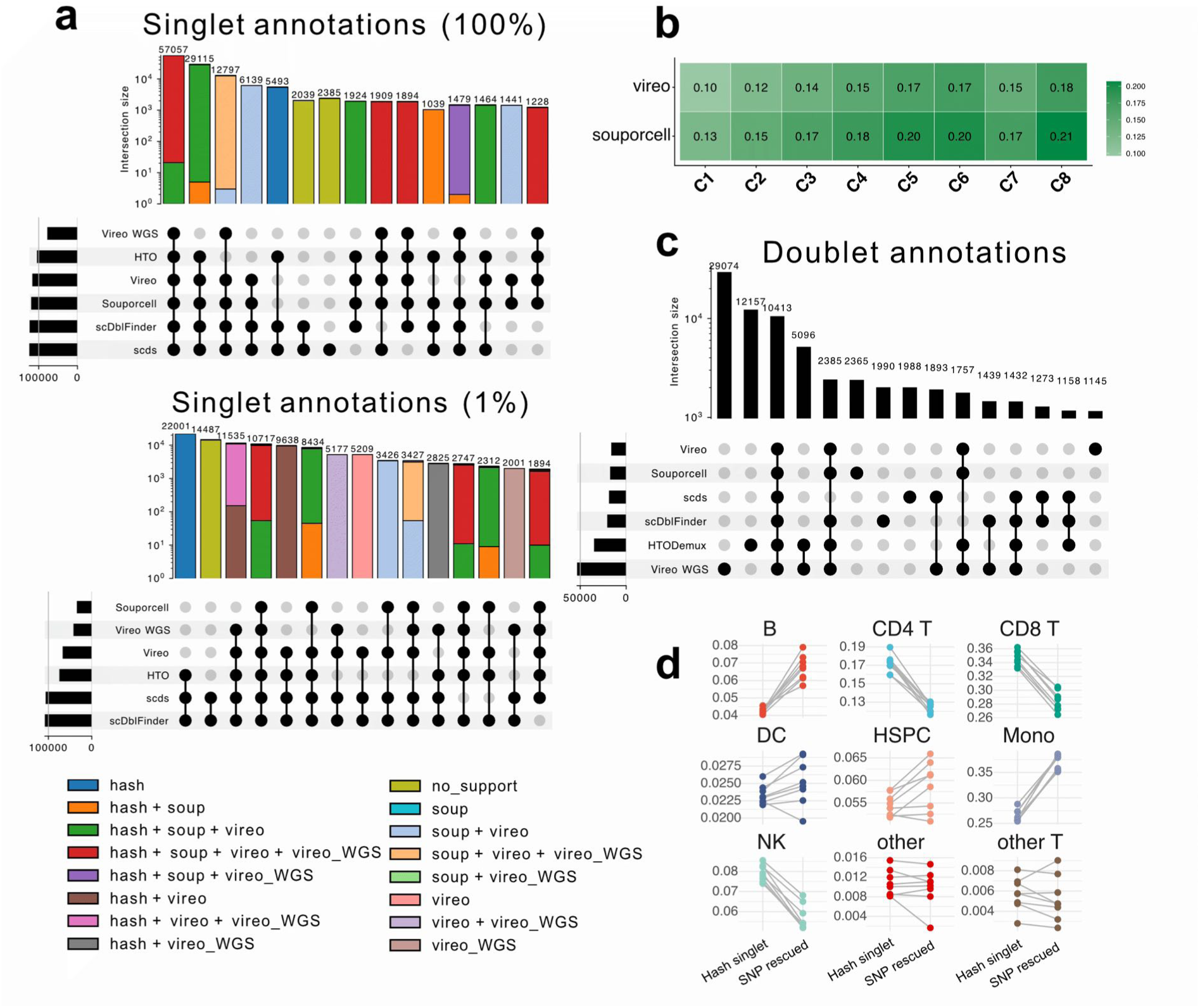
Concordance of demultiplexing and doublet detection strategies in assigning true singlets. Singlets and doublets were annotated using a combined approach that incorporated demultiplexing-based identification of donor-resolved singlets and cross-sample multiplets, together with two transcriptome-based doublet detection methods. Demultiplexing was performed using HTODemux (HTO-based demultiplexing), Vireo and Souporcell (SNP-based demultiplexing using a reference SNP panel from the 1000 Genomes Project), and Vireo WGS (SNP-based demultiplexing using a donor-specific SNP panel derived from whole-genome sequencing). Transcriptome-based doublet detection was carried out using scDblFinder and scds. **(a)** UpSet plot (top) showing the overlap of cells annotated as singlets across the six methods. Bars indicate the number of cells classified as singlets by the intersection between the specified methods; only cells with valid patient assignments from the Cell Ranger-called barcode set are included in this analysis. The stacked categories filled in the bar plots represent the method combinations that agreed on the same donor for which the majority voting succeeded. Cells in the top 15 most frequent intersections are plotted. The sets represented constitute 92.7% of the total cells. The bottom UpSet plot shows the exact visualization using the downsampled 1% dataset; cells in the top 15 most frequent intersections represent 74.8% of the total cells. **(b)** Fraction of additional singlet assignments recovered by SNP-based demultiplexing relative to HTO-only demultiplexed singlets across channels. **(c)** Same as (a) but for cells annotated as doublets. Cells in the top 15 most frequent intersections are plotted. The sets represented constitute the 87.1% of the total cells. **(d)** Paired comparison of cell-type fractions between HTO-defined singlets and SNP-rescuable HTO-doublets across channels. Points represent individual channels; lines connect paired measurements from the same channel. See also **Supplementary Figs. S2-S5.**

To assess how sequencing depth affects demultiplexing performance, we applied a read downsampling approach on the multiplexed CITE-seq data at 50% (ds50), 10% (ds10), and 1% (ds01) of the original sequencing depth. Subsequently, all demultiplexing methods were run on these datasets. Downsampling led to a systematic decrease in total UMI counts, ADT counts, and the number of unique detected features across all channels (**Supplementary Fig. S1a-d**). The number of informative SNPs recovered per channel dropped sharply with downsampling for both cellsnp-lite + Vireo and Souporcell, with a more pronounced loss for Souporcell at lower depths (**Supplementary Fig. S1e**). As a result, SNP-based singlet recovery declined substantially with reduced SNP recovery (**Supplementary Fig. S1f**). Assignment trajectories confirmed that most singlets remained stable at ds50, while very low coverage at ds01 shifted a large fraction of cells into the unassigned category (**Supplementary Fig. S1f**). Hash-based demultiplexing was less sensitive to reduced sequencing depth than SNP-based demultiplexing methods (**Supplementary Fig. S1f**). At the ds01 level, only hash-recoverable donors remained as the most frequent singlet category (**Fig. 3a**). In the original full-depth dataset, Vireo and Souporcell reported higher singlet fractions than HTODemux across all channels, indicating that this relative behavior is reproducible and not driven by channel-specific artifacts. Souporcell recovered slightly more singlets across all channels in the full-depth dataset, reflecting small method-specific performance variability in both contexts (**Fig. 3b**). These results demonstrate that SNP-based demultiplexing recovers additional singlets at full sequencing depth but is disproportionately sensitive to reduced coverage, supporting the recommendation to combine both strategies.

### Combining orthogonal demultiplexing methods rescues misclassified singlets and reduces cell-type-dependent bias

We next examined agreement on doublet annotations across methods. The UpSet plot revealed that hash-only doublets (cells labeled solely by HTODemux) formed the second largest single-method doublet group after Vireo WGS. In contrast, consensus doublets identified by all six methods formed a smaller intersection (**Fig. 3c**). This raised the question of whether hash-only doublets are true multiplets or false positives resolvable with orthogonal evidence.

Hash-only doublets were enriched in specific cell types such as B cells and monocytes, indicating that false-positive hashing calls enrich in populations with particular surface markers or transcriptional profiles (**Supplementary Fig. S2e**). SNP-rescuable hash doublets (called as doublets by HTODemux but as singlets by both Vireo and Souporcell) had UMI counts and gene distributions similar to consensus singlets (**Supplementary Fig. S2f-g**), supporting their status as false-positive hash calls. Reclassifying these cells altered cell-type composition: SNP-rescuable hash doublets had different cell-type proportions than HTO-defined singlets, with certain populations disproportionately affected (**Fig. 3d**).

Collectively, these results reveal that disagreement between demultiplexing strategies is not randomly distributed across cells. Instead, hash-only doublets concentrate in specific cell types and quality strata, indicating that the pattern of cross-method discordance carries structured information about cell identity and data quality. This observation transforms what is conventionally treated as a method disagreement requiring resolution into a diagnostic signal that can guide cell selection. Building on this insight, we formalized the concordance patterns into a decision framework that uses method agreement and disagreement to stratify cells by classification confidence.

### A concordance-based framework uses method-disagreement to stratify cell quality

Based on the systematic method-level comparisons described above, we devised a concordance-based decision framework for high-confidence singlet selection (**Fig. 4**). Compared to other established frameworks, Split-flow employs a staged strategy. We first classify cells as singlets, unassigned, or doublets using a concordance-based decision framework. This classification is then integrated with transcriptome-based multiplet detection results and donor assignment agreement to produce five final categories: (i) high-confidence singlets, defined as cells with concordant singlet classification across demultiplexing methods and consistent donor assignment; (ii) low-confidence singlets, with majority singlet support but weaker or partial donor evidence; (iii) doublets, concordantly classified as multiplets or flagged by transcriptome-based detection; (iv) unassigned, for which no majority classification could be achieved; and (v) donor-assignable but unrecoverable cells, for which donor-related evidence exists but is contradictory across methods, preventing majority voting on the donor level (see Methods and Supplementary Table).

**Figure 4.**
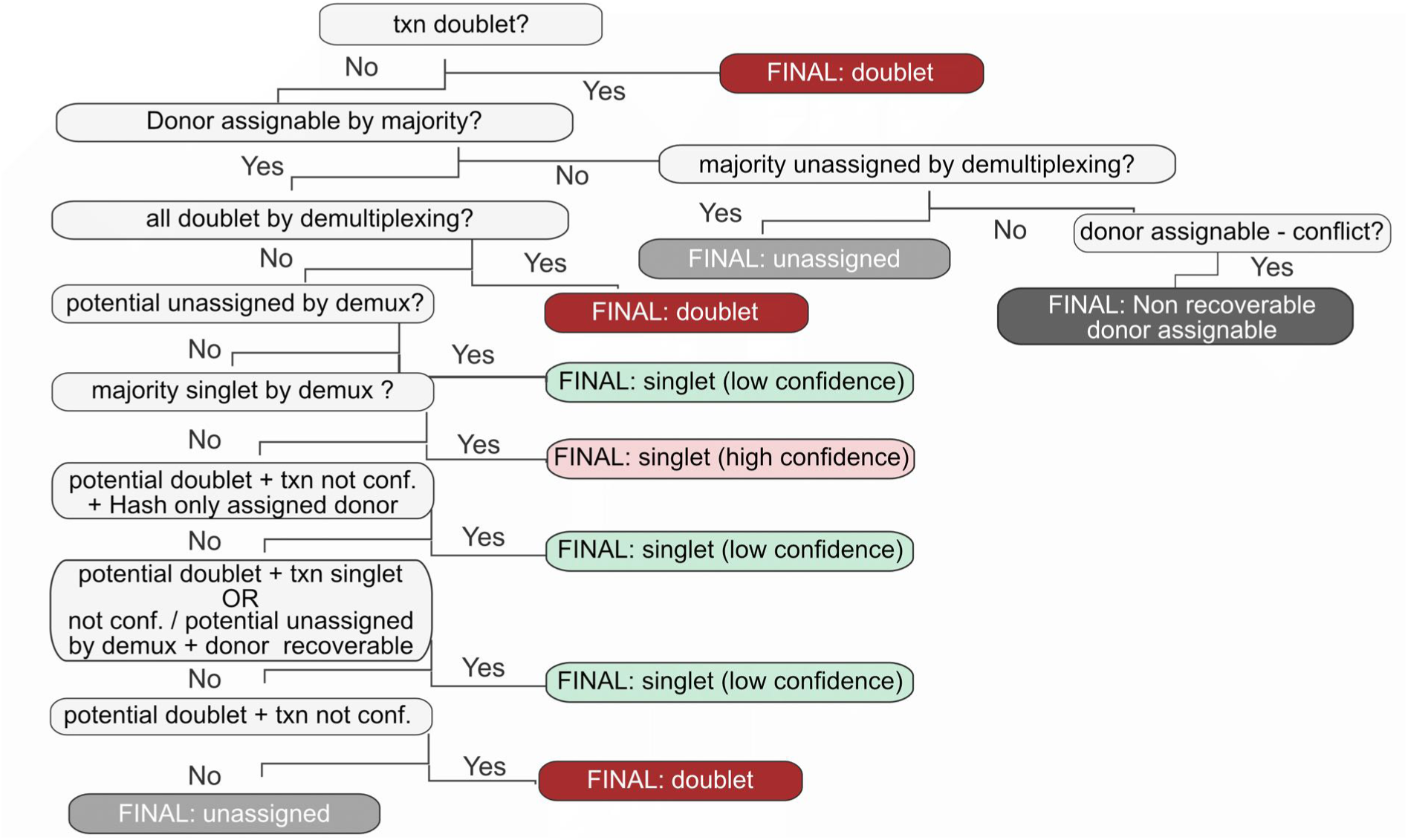
Decision-making framework for demultiplexing and high-confidence cell selection. Concordance-based framework for demultiplexing and high-confidence cell selection. Cell calling defines the initial set of cell-containing droplets, which are then subjected to parallel demultiplexing and doublet detection alongside quality evaluation. Outputs are integrated in a post hoc concordance evaluation that informs terminal exclusion of consensus doublets and consensus unassigned cells. The remaining high-confidence singlets are routed to unimodal or multimodal feature spaces for downstream analysis.

Before assignment, the Split-flow framework retains intermediate concordance categories. These include, for example, cells classified as singlets by SNP-based methods but as doublets by hashing, or cells lacking a majority vote in initial singlet annotation, which we define and label as “not confident.” This approach enables users to evaluate borderline cells with explicit confidence annotations rather than forcing binary decisions. By retaining intermediate concordance categories, the framework preserves information that single-tool selection or majority voting would discard. To verify whether this stratification accurately reflects quality differences, we analyzed QC metrics and embedding patterns across the final categories.

### Concordance categories reflect genuine quality gradients, not random noise

Using our concordance-based decision framework, we assigned cells to five final classification categories, derived from an expanded set of 33 intermediate classes that captured combinations of demultiplexing, doublet detection, and donor assignment outcomes. When we assessed whether these categories reflected biologically meaningful differences, the intermediate classes showed a gradient of quality. High-confidence singlets had higher total RNA and ADT counts and more detected genes than low-confidence singlets (**Supplementary Fig. S3a, b**). In contrast, mitochondrial fraction and ambient RNA contamination, estimated per cell by total count correction with CellBender, increased steadily toward lower-confidence or unassigned classes (**Supplementary Fig. S3c-e**). These quality differences were also evident in the UMAP embedding, where final classification categories were distributed non-randomly, with low-confidence or unassigned cells localizing to specific regions on the UMAP (**Fig. 5a-b**). Cells with high ambient RNA levels and those with higher mitochondrial content clustered together in regions enriched for discordant and unassigned cells (**Fig. 5c-d**).

**Figure 5.**
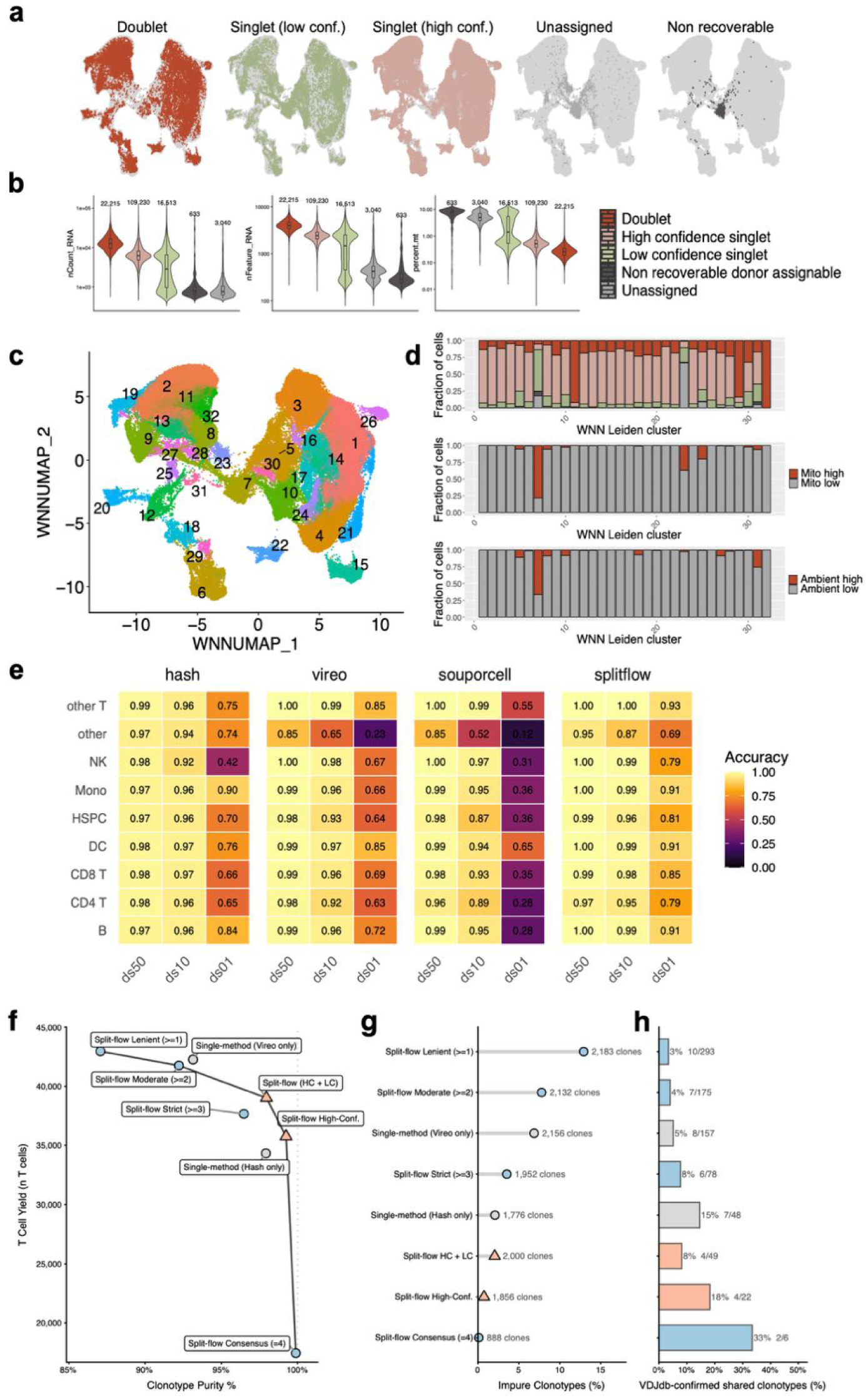
Application of the Split-flow workflow to identify quality differences, artifact-enriched cell populations and T-cell receptor concordance. **(a)** WNN UMAP colored by final cell category assignments. **(b)** Distribution of QC metrics per cell across final assignment categories. **(c)** WNN UMAP colored by Leiden clusters (resolution = 0.4) with cluster labels displayed. **(d)** Fraction of cells colored by final category, mitochondrial content, and high ambient RNA status across WNN Leiden clusters. **(e)** Heatmap showing the accuracy of donor assignment for singlets across annotated bone marrow cell types following the downsampling trajectory (ds50, ds10, and ds01). Accuracy was determined relative to donor assignment from the full dataset and assessed separately for each method. **(f)** Scatter plot showing the T cell yield vs clonotype purity for different demultiplexing strategies. **(g)** Lollipop chart showing the percentage of impure (found in more than one donor) clonotypes across strategies, with absolute clone counts indicated. **(h)** Horizontal bar chart showing the precision of shared clonotype identification as given by VDJdb CDR3 beta overlaps. Absolute number of confirmed vs total shared clones is shown.

### Split-flow mitigates depth-dependent biases in cell-type recovery

We next evaluated how assignment accuracy changed across cell types and downsampling levels for different demultiplexing strategies. Assignment accuracy varied by cell type, demultiplexing strategy, and sequencing depth, indicating that the method choice can impact which biological populations are preserved under reduced coverage (**Fig. 5e**). Notably, singlets identified by the Split-flow framework were less affected by depth-related sensitivity loss than those identified through hash-based or SNP-based demultiplexing strategies alone (**Fig. 5e**). Overall, these findings demonstrate that the concordance-based classification in Split-flow captures genuine quality gradients, and that its final categorization, compared to single demultiplexing methods, helps mitigate the effects of coverage-related biases that can cause the loss of biologically relevant populations. However, all analyses thus far have relied on computational QC metrics to evaluate concordance categories. To establish that these categories reflect genuine biological quality differences rather than technical artifacts of the metrics themselves, an independent biological validation is needed.

### Concordance-based selection maximizes biological accuracy of TCR recovery

To independently validate that concordance categories reflect genuine biological quality differences, we leveraged T cell receptor (TCR) sequences as an orthogonal readout. Because productive TCR rearrangements are somatically unique, clonotypes shared across donors in a multiplexed pool indicate demultiplexing error unless they correspond to known public TCR sequences. We quantified T cell yield, clonotype purity, and the biological plausibility of cross-donor sharing across demultiplexing strategies. Yield and purity showed a pronounced trade-off (**Fig. 5f**). Lenient consensus (assignment by ≥1 method) recovered the most T cells (42,970) and clonotypes (2,183) but with 12.9% cross-donor contamination (**Fig. 5g**).

Single-method approaches showed intermediate performances: hash-only yielded 34,325 T cells at 97.9% purity, Vireo-only recovered more cells (42,276) at lower purity (93.1%). Split-flow High-Confidence achieved 99.2% purity while retaining 34,756 T cells, and the HC+LC stratum provided a practical balance at 98% purity with 39,020 T cells (**Fig. 5f-g**).

To assess whether cross-donor clonotypes represent genuine public TCRs or demultiplexing artifacts, we cross-referenced shared clonotypes against VDJdb, a curated database of validated TCR sequences [37, 38]. Lenient strategies yielded the most shared clonotypes (293) but the lowest VDJdb confirmation rate (3.4%). Split-flow High-Confidence achieved 18.2% precision (4/22 confirmed), a 5.3-fold enrichment over the lenient baseline, while the Consensus stratum reached 33.3% (2/6) (**Fig. 5h**). All enrichments over the private-clonotype background rate (∼0.7%) were significant (Fisher’s exact test, all P<0.001). These results demonstrate that concordance categories capture genuine differences in biological signal fidelity: stricter concordance requirements progressively filter out demultiplexing artifacts while enriching for biologically plausible TCR sharing events.

### Split-flow recovers high-quality singlets from AML snMultiome-seq despite low hashing efficiency

The preceding analyses established that concordance-based preprocessing captures genuine quality differences in multiplexed CITE-seq data from MM patients, validated both by computational QC metrics and by independent biological evidence from TCR sequences. A key remaining question is whether this framework transfers to different disease contexts and assay types. To address this, we applied Split-flow to AML patient samples profiled by snMultiome-seq. Two independently processed pools, each comprising samples from four AML patients, were loaded onto 10x Chromium channels with approximately 20,000 nuclei per channel (**Fig. 6**). In this assay, RNA and ATAC readouts were acquired simultaneously from the same nuclei, and sample barcoding was performed using CMOs, consistent with the strategy outlined in **Table 1**.

**Figure 6.**
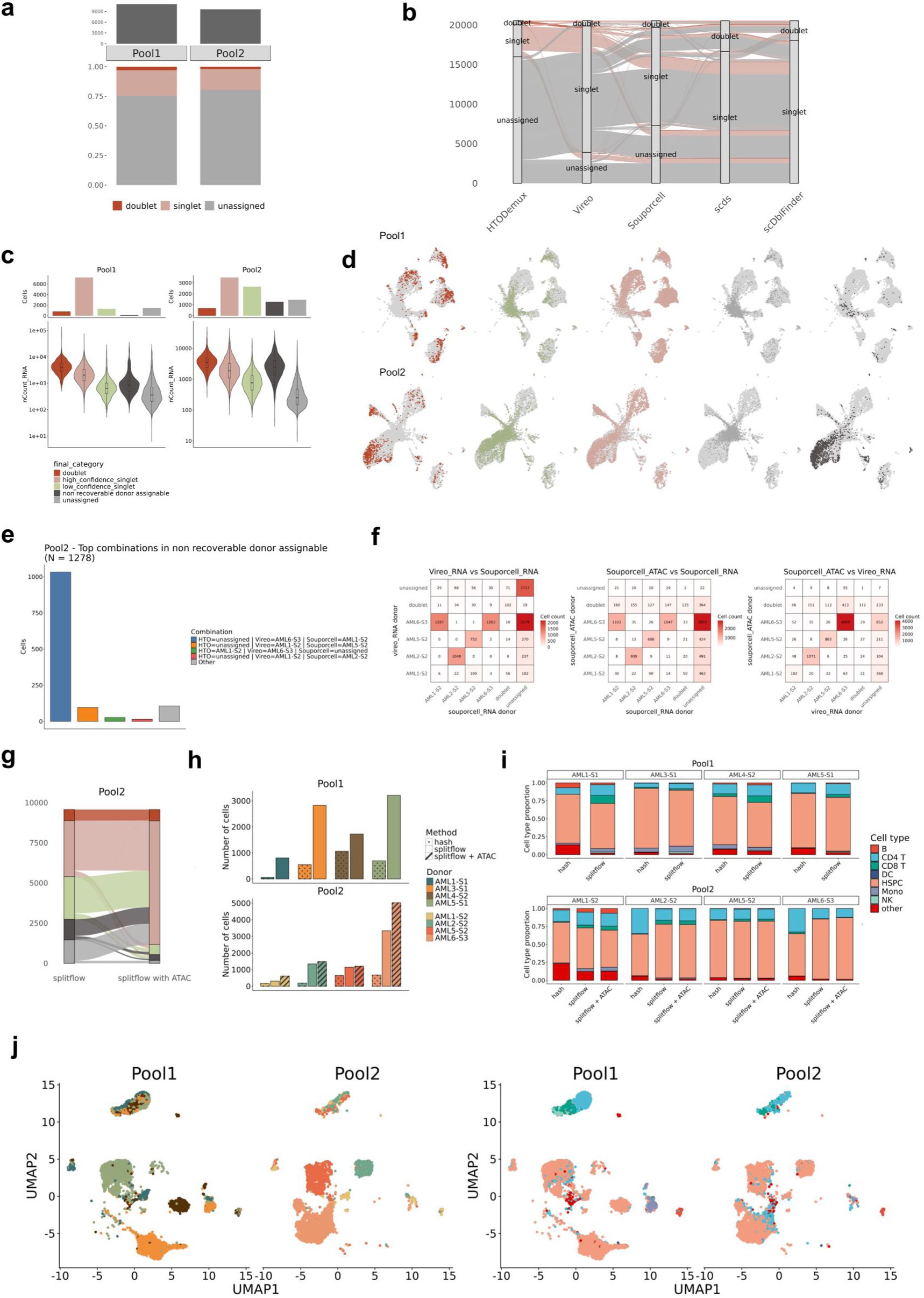
Assessment of the Split-flow workflow in AML Multiome data. **(a)** Stacked bar plots showing the fraction of cells classified as singlets, doublets, or unassigned by CMO-based demultiplexing (HTODemux) for each AML pool. Bars above indicate the total number of Cell Ranger–called cells per pool. Each pool represents one 10x Chromium channel containing four multiplexed AML samples (∼20,000 cells loaded). **(b)** Alluvial plot comparing cell assignment outcomes across demultiplexing and doublet detection methods. Flows represent cells transitioning between classification categories (singlet, doublet, unassigned) across HTODemux, Vireo, Souporcell, scds, and scDblFinder. Flow widths correspond to cell counts, and colors indicate HTODemux classifications as defined in (a). **(c)** Bar plots showing the total number of cells assigned to each final classification category for each pool based on the Split-flow decision framework (Fig. 4) (top), and violin plots showing the distribution of UMI counts (nCount_RNA) per cell across these categories for each pool (bottom). **(d)** UMAP embedding colored by final classification category, shown separately for each pool. **(e)** Bar plot showing the top four combinations of donor assignments across HTODemux, Vireo, and Souporcell for cells classified as non-recoverable donor assignable in Pool2. **(f)** Heatmaps showing agreement in donor assignment across demultiplexing methods. Pairwise comparisons between Vireo (RNA) and Souporcell (RNA) (left), Souporcell (ATAC) and Souporcell (RNA) (middle), and Souporcell (ATAC) and Vireo (RNA) (right). Values indicate cell counts. **(g)** Alluvial plot comparing final cell classification categories between two Split-flow configurations for Pool2: one including Souporcell (RNA) and one including Souporcell (ATAC) for demultiplexing. Flows represent cell transitions between categories. **(h)** Bar plots showing the number of cells recovered per donor across methods. Pool1 (top) compares HTODemux (hash) and Split-flow, while Pool2 (bottom) additionally includes Split-flow using Souporcell (ATAC) instead of Souporcell (RNA). For Split-flow, only cells classified as high- or low-confidence singlets were included. **(i)** Stacked bar plots showing cell-type composition per donor among cells recovered across methods. Pool1 (top) compares HTODemux (hash) and Split-flow, while Pool2 (bottom) additionally includes Split-flow using Souporcell (ATAC) instead of Souporcell (RNA). Cell types are based on level 1 annotations obtained using Azimuth with a human bone marrow reference. Only cells with nCount_RNA > 500 were included. For Split-flow, only cells classified as high- or low-confidence singlets were retained. **(j)** WNN UMAP (RNA and ATAC) of high- and low-confidence singlets identified by Split-flow for Pool1 and by Split-flow using Souporcell (ATAC) for Pool2, shown separately by pool and colored by donor (left) and cell type (right). Only cells with nCount_RNA ≥ 500 and nCount_ATAC ≥ 1000 were included. Batch effects between pools were corrected using Harmony prior to integration.

CMO-based demultiplexing using HTODemux yielded low singlet fractions across both AML multiome pools, suggesting limited hashing efficiency (**Fig. 6a**). Consistently, CLR-normalized CMO signals showed poor separation between positive and background populations. Distributions across donors were broad and overlapping, indicating a low signal-to-noise ratio that likely impairs accurate hash-based classification (**Supplementary Fig. S4a**). Applying Split-flow to the RNA modality enabled systematic comparison of cell classifications across methods. The alluvial plot in **Fig. 6b** highlights substantial discrepancies in cell classification. Many cells unassigned by CMO-based demultiplexing were reassigned as singlets by SNP-based approaches, underscoring the benefit of combining orthogonal strategies.

The concordance-based decision framework classified cells into five final categories from 27 intermediate classes (**Fig. 6c; Supplementary Fig. S4b**). In Pool1, these categories followed a clear quality gradient, with decreasing RNA content (**Fig. 6c**) and detected gene counts (**Supplementary Fig. S4c**) from high-confidence singlets to unassigned cells. In contrast, this trend was less evident in Pool2, where non-recoverable donor-assignable cells exhibited relatively higher RNA content and constituted a larger fraction of the dataset. These patterns were reflected in the UMAP embeddings (**Fig. 6d**), where lower-confidence categories in Pool1 were predominantly localized to intermediate regions of the embedding, consistent with reduced data quality, whereas in Pool2 they were more broadly distributed and overlapped with regions occupied by high-confidence singlets.

### ATAC-based demultiplexing resolves donor assignment discordance in snMultiome-seq

Given the large fraction and high RNA content of non-recoverable donor-assignable cells in Pool2 (**Fig. 6c**), we hypothesized that these represent high-quality cells with discordant donor assignments. We therefore examined the most frequent assignment combinations from HTODemux, Vireo, and Souporcell within this category (**Fig. 6e**). While many cells were unassigned by HTODemux, consistent with low hashing efficiency, the dominant pattern was discordance between genotype-based methods, with Vireo and Souporcell frequently assigning different donors to the same cells. To investigate this discordance within the Split-flow framework, which integrates Vireo and Souporcell for RNA, we additionally leveraged the ATAC modality by performing Souporcell-based demultiplexing on ATAC and comparing these assignments to those from RNA (**Fig. 6f**). Substantial discordance was observed between Vireo and Souporcell based on RNA, as well as between Souporcell assignments derived from RNA and ATAC. In contrast, Souporcell for ATAC showed improved concordance with Vireo. Based on these observations, we re-ran the Split-flow classification for Pool2 by replacing RNA with ATAC for Souporcell. This led to substantial reassignment of cells across categories, with many previously non-recoverable donor-assignable and unassigned cells transitioning into singlet categories (**Fig. 6g; Supplementary Fig. S4d**). This redistribution was accompanied by reduced RNA and gene counts in non-recoverable donor-assignable cells (**Supplementary Fig. S4e**). Lower-confidence categories were predominantly localized to intermediate regions of the embedding, consistent with reduced data quality (**Supplementary Fig. S4f**).

### Split-flow increases donor recovery and alters inferred cell-type composition

To quantify donor recovery, we compared the number of cells assigned per donor across methods (**Fig. 6h**). CMO-based demultiplexing alone yielded substantially fewer cells per donor. In contrast, the Split-flow framework markedly increased recovery, with ATAC-based demultiplexing in Pool2 providing additional gains.

Comparing cell-type composition across methods, Split-flow altered the relative proportions of cell types recovered per donor, indicating that low hashing efficiency can bias biological conclusions (**Fig. 6i**). ATAC-based demultiplexing in Pool2 had minimal impact on cell-type composition, suggesting that this refinement primarily improves donor assignment without altering biological distributions. Finally, we applied a WNN UMAP embedding based on both RNA and ATAC modalities (**Fig. 6j**). The integrated embedding revealed both shared and donor-specific structure across pools. Immune cell populations, including T cells, formed well-mixed clusters across donors, consistent with shared microenvironmental cell states. In contrast, progenitor-like populations (HSPCs) formed more distinct, donor-enriched clusters, suggesting inter-patient heterogeneity consistent with malignant cell populations, likely representing leukemic blasts. Together, these results demonstrate that the Split-flow framework enables recovery of high-quality cells suitable for downstream multimodal analysis, allowing both shared and patient-specific biological features to be resolved.

### The Split-flow framework generalizes to independently profiled AML scRNA-seq and scATAC-seq

To further assess whether the Split-flow framework generalizes across assay types, we applied it to AML patient samples profiled using independent scRNA-seq and scATAC-seq experiments. Three pools, each comprising samples from three AML patients, were loaded onto 10x Chromium channels (∼20,000 cells per channel). Sample multiplexing was performed using CMOs for scRNA-seq and Tn5-based barcoding for scATAC-seq (**Table 1**).

CMO-based demultiplexing of the scRNA-seq data yielded moderate singlet fractions across all three pools (**Fig. 7a**), consistent with limited hashing efficiency. This was further supported by poor separation of CLR-normalized CMO signals across donors (**Supplementary Fig. S5a**).

**Figure 7.**
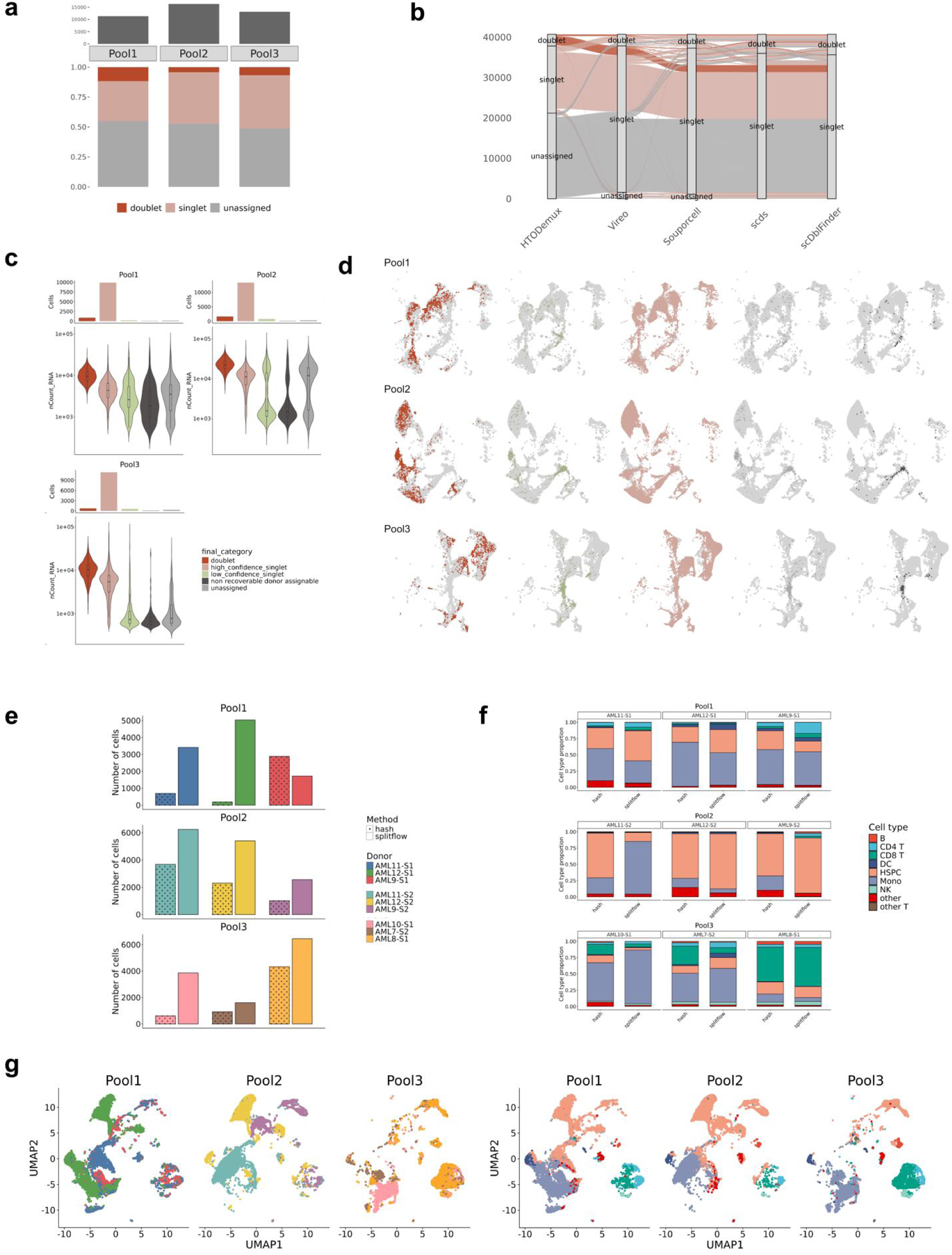
Split-flow workflow in AML scRNA-seq and scATAC-seq data. **(a)** Stacked bar plots summarizing the proportions of cells classified as singlets, doublets, or unassigned by CMO-based demultiplexing (HTODemux) across AML pools. Bars above indicate the total number of Cell Ranger–detected cells per pool. Each pool corresponds to a single 10x Chromium channel containing three multiplexed AML samples (∼20,000 cells loaded). **(b)** Alluvial plot illustrating concordance and discordance in cell assignments across demultiplexing and doublet detection methods. Flows represent transitions of individual cells between classification states (singlet, doublet, unassigned) across HTODemux, Vireo, Souporcell, scds, and scDblFinder. Flow widths reflect cell counts, and colors denote HTODemux classifications as defined in (a). **(c)** Bar plots showing the number of cells assigned to each final classification category per pool according to the Split-flow decision framework (Fig. 4) (top), alongside violin plots depicting per-cell UMI count distributions (nCount_RNA) across these categories (bottom). **(d)** UMAP embedding colored by final classification category, displayed separately for each pool. **(e)** Bar plots showing the number of cells recovered per donor across methods for each pool. For Split-flow, only cells classified as high- or low-confidence singlets are included. **(f)** Stacked bar plots showing the distribution of cell types per donor among recovered cells across methods. Cell types were assigned using Azimuth level 1 annotations with a human bone marrow reference. Only cells with nCount_RNA > 1000 were included, and for Split-flow, only high- and low-confidence singlets were retained. **(g)** UMAP embedding of high- and low-confidence singlets identified by Split-flow, shown separately for each pool and colored by donor identity (left) and cell type annotation (right). Only cells with nCount_RNA ≥ 1000 were included. Batch effects between pools were corrected using Harmony.

Integration of SNP-based approaches within the Split-flow framework improved cell classification, with the alluvial plot revealing substantial reassignment of cells from unassigned or doublet categories to singlets (**Fig. 7b**). The concordance-based framework classified cells into five categories with consistent distributions across pools (**Fig. 7c; Supplementary Fig. S5b**), again showing a quality gradient from high-confidence singlets to lower-confidence categories (**Fig. 7c; Supplementary Fig. S5c**). This pattern was reflected in UMAP embeddings, where lower-confidence cells were predominantly localized to intermediate regions of the embedding, consistent with reduced data quality (**Fig. 7d)**. Split-flow substantially increased the number of cells recovered per donor compared to CMO-based demultiplexing alone (**Fig. 7e**). As in the multiome data, Split-flow altered the relative proportions of cell types recovered per donor, indicating that reliance on hashing alone can bias inferred cellular composition (**Fig. 7f**). Finally, the UMAP visualization of high- and low-confidence singlets demonstrated coherent biological structure across pools, with both shared and donor-specific patterns evident (**Fig. 7g**).

In parallel, scATAC-seq samples were multiplexed using Tn5-based barcoding, which introduces sample identity during library construction and bypasses CMO labeling efficiency limitations. This resulted in balanced donor representation and consistent quality metrics across pools, with clear donor separation in the embedding (**Supplementary Fig. S6a-d**), indicating robust sample assignment independent of computational demultiplexing. Together, these results confirm that the Split-flow framework generalizes across assay types and enables robust donor assignment and cell recovery in independently profiled scRNA-seq and scATAC-seq datasets.

## Discussion

Preprocessing multiplexed single-cell sequencing data requires multiple analytical steps, the outcomes of which depend on the type and quality of the input samples, the available sequencing readouts, the combination of computational methods, and the criteria used for cell selection. Suboptimal preprocessing decisions can systematically exclude biologically relevant cell populations while retaining low-quality cells, directly biasing downstream biological interpretation. Here, we show that disagreement between orthogonal preprocessing methods is not merely noise to be resolved but a structured, interpretable signal of cell quality. By retaining intermediate concordance categories and stratifying cells by the pattern of cross-method agreement, Split-flow uses method discordance as a diagnostic tool to identify cells for downstream analysis.

Existing approaches to demultiplexing apply three main strategies. (i) Specific assignment methods are optimized [21, 39–42]. (ii) Benchmarking and consensus studies compare tools and propose combination strategies. These include Demuxafy’s decision tree for tool selection [43], Mylka et al.’s comparison of multiplexing strategies [30], hybrid approaches [44], and ensemble methods such as Ensemblex [45] and scConSensus [46], and an independent benchmark of computational demultiplexing methods on snRNA data [47]. (iii) Integrated pipelines provide end-to-end workflows: hadge selects the best-performing tool per pool [22], YASCP provides a modular pipeline for population genetics [23], and CellDemux performs coherent genetic demultiplexing across sequencing modalities.

Split-flow differs from these approaches in how it treats cross-method disagreement. Current approaches select the best tool (hadge), apply majority voting (Ensemblex, scConSensus), or focus on within-tool coherence (CellDemux). In contrast, Split-flow retains intermediate concordance categories and treats the pattern of disagreement as a quality signal. It preserves borderline cells with explicit confidence labels rather than forcing binary inclusion or exclusion. Beyond this conceptual distinction, our study addressed three practical gaps. First, Split-flow was validated on real clinical samples from hematopoietic malignancies with variable viability, high ambient RNA, and immunophenotypic shifts, rather than on healthy-donor PBMCs or simulated data. Second, its experimental design was linked to computational preprocessing by systematically evaluating barcoding strategies. Third, we implemented a “run everything, decide later” architecture that defers terminal classification to post hoc concordance evaluation, preserving information that early tool selection or majority voting would discard (**Fig. 3**, **Fig. 5e**).

The multiplexed CITE-seq experiment demonstrated high technical reproducibility across channels, providing a reliable baseline for evaluating concordance-based classification (**Fig. 2a-c**). Discordant cells cluster within specific cell types and quality strata (**Fig. 5**), and rescued cells exhibited QC metrics comparable to those of concordant singlets (**Supplementary Fig. S2f,g**). These findings provide empirical justification for the concordance-based approach over single-tool or majority-vote strategies.

The differential sensitivity of demultiplexing strategies to sequencing depth has direct implications. SNP-based methods lose informative variants more rapidly than hash-based approaches under downsampling, with Souporcell declining more steeply than Vireo (**Supplementary Fig. S1a-f**). We conclude that researchers should combine genetic and hash-based demultiplexing when possible. The transferability from MM CITE-seq to AML samples across snMultiome-seq and separate scRNA-seq/scATAC-seq (**Fig. 6-7**) demonstrates that our framework generalizes across hematopoietic malignancies and assay types while remaining sensitive to disease-specific sample properties.

The TCR-based validation introduced in our study provides independent biological evidence that concordance categories capture genuine quality differences beyond QC metric gradients. Because productive TCR rearrangements are somatically unique, cross-donor clonotype sharing is an unambiguous indicator of demultiplexing error unless the shared sequences correspond to known public TCRs. The 5.3-fold enrichment of VDJdb-confirmed sequences among Split-flow High-Confidence shared clonotypes (18.2% vs 3.4% precision) demonstrates that concordance-based classification preserves biological truth at the level of individual immune receptor sequences. This is particularly relevant for clonal tracking and disease-associated TCR signatures, where cross-donor contamination can mimic convergent selection. The HC+LC stratum achieves 98% clonotype purity while retaining 92% of detected clonotypes, suggesting a practical operating point for studies requiring both high recovery and biological accuracy.

ATAC-based genetic demultiplexing can resolve donor-assignment conflicts when RNA-based methods disagree. In Pool2, replacing RNA with ATAC for Souporcell increased agreement and recovered otherwise unassigned cells (**Fig. 6e-g**). This demonstrates Split-flow’s modularity: because evidence generation is separate from classification, modality-specific inputs can be substituted without restructuring the pipeline. This capability is conceptually related to CellDemux [46]; our Pool2 analysis independently demonstrates that cross-modal demultiplexing can substantially improve cell recovery. A complementary multiplexing approach is presented by MULTI-ATAC, which uses pooled transposition with sample-specific Tn5 transposomes [48]. The same principle extends upstream: default multiome cell calling with CellRanger-arc has been reported to miss over half of nuclei and to be biased against specific cell types compared with a modality-aware statistical alternative [17], consistent with our view that preprocessing bias is not limited to demultiplexing.

Several limitations should be noted. First, applicability to solid tumors or non-malignant tissues with different cellular compositions, RNA quality, or ambient background remains to be evaluated. Second, concordance thresholds are heuristic decision rules rather than formally optimized cutoffs, though monotonic quality gradients across categories support their relevance (**Fig. 5, Supplementary Fig. S3**). Third, full ground-truth genotype panels were not available for all datasets, and the framework has not been tested with closely related donors, where SNP-based demultiplexing is expected to be less informative. Finally, we evaluated a specific tool set (HTODemux, Souporcell, Vireo, scds, and scDblFinder), and extension to other methods may require adapting the concordance rules.

A key strength of our study is the combination of computational QC validation with independent biological evidence from TCR sequences. To our knowledge, using clonotype-level cross-donor sharing as a metric for preprocessing accuracy has not been applied previously in this context. The 5.3-fold enrichment of VDJdb-confirmed public TCR sequences in the high-confidence stratum provides direct evidence that concordance-based classification preserves biological signal fidelity at the level of individual immune receptor sequences. This approach could serve as a general validation strategy for preprocessing pipelines applied to immune-rich tissues. More broadly, the principle that structured disagreement between analytical methods carries diagnostic information may extend beyond single-cell preprocessing. Analogous concordance-based strategies could be applied wherever multiple computational approaches address the same classification task on complex biological data.

## Conclusions

We conclude that disagreements among orthogonal preprocessing methods in multiplexed single-cell experiments should be treated as a structured signal reflecting genuine differences in cell quality rather than noise to be minimized. Split-flow operationalizes this principle through a concordance-based decision framework that retains intermediate quality categories with explicit confidence annotations, enabling transparent evaluation of borderline cells. Independent validation using T cell receptor sequences confirms that concordance categories capture biological signal fidelity. The framework generalizes across hematopoietic malignancies and assay types, with multi-assay validation across CITE-seq, snMultiome-seq, and separate scRNA-seq/scATAC-seq within a single study. The framework’s modularity further enables cross-modal demultiplexing, as demonstrated by using ATAC-based genotyping to resolve donor-assignment discordance in snMultiome-seq data. Implemented as an open-source Nextflow DSL2 pipeline with dual R/Seurat and Python/Scanpy output, Split-flow provides a practical, reproducible framework for linking experimental design to data preprocessing in multiplexed single-cell studies.

## Methods

### CITE-seq of MM samples

Viably frozen BMMCs from MM patients were processed for CITE-seq as described elsewhere. Individual patient samples were multiplexed by labeling cells with 0.1 µg of TotalSeq anti-human Hashtag oligonucleotide antibodies (Hashtag 11–24; BioLegend). Subsequently, cells were incubated for 30 minutes at 4 °C with a TotalSeq™-C antibody cocktail (BioLegend) to enable simultaneous detection of cell surface proteins. Immunoprofiling was performed with the Chromium Next GEM Single Cell 5ʹ Reagent Kit v2 (Dual Index) according to the user guide, which features barcode technology for cell surface protein & immune receptor mapping (10x Genomics; CG000330 Rev A).

### scMultiome-seq of AML samples

For scMultiome-seq of AML samples, samples were thawed and processed without prior viable cell sorting. Nuclei were isolated using ice-cold nuclei isolation buffer, incubated briefly on ice, and pelleted using a swing-bucket centrifuge to minimize nuclei loss. Isolated nuclei were resuspended in PBS supplemented with RNase inhibitor, counted, and adjusted to the desired concentration. For sample multiplexing, approximately 500,000 nuclei per sample were labeled with CMOs, followed by incubation on ice and sequential wash steps to remove excess oligos. After the final wash, nuclei were resuspended in 1× Multiome nuclei buffer, quantified, and pooled in equal numbers, with four samples multiplexed per pool. Further processing was with the Chromium Next GEM Single Cell Multiome ATAC and Gene Expression kit (10x Genomics). For oligo sequences and CMO labeling protocol details, see [49].

### Separate scRNA-seq and snATAC-seq of AML sample

For separate scRNA-seq and snATAC-seq of AML samples, cryopreserved cells were thawed, washed in PBS supplemented with 1% FCS, and counted to assess viability. Cells were stained with CD34 and CD117 antibodies following Fc receptor blocking and sorted by FACS to enrich for target populations. Sorted cells were collected in FACS buffer and subsequently quantified. For scRNA-seq, approximately 500,000 cells per sample were labeled with CMOs, followed by incubation and wash steps to remove excess oligos. After hashing, cells were processed using the Chromium Next GEM Single Cell 5’ Gene Expression kit (10x Genomics) according to the manufacturer’s instructions. For snATAC-seq, remaining cells were used for nuclei isolation by incubation in ice-cold nuclei isolation buffer, followed by centrifugation and resuspension in nuclei buffer. Nuclei were counted, and indexed Tn5 transposase complexes were assembled using pre-annealed oligonucleotides. Approximately 50,000–180,000 nuclei per reaction were subjected to tagmentation at 37 °C, followed by washing, pooling in equal proportions, and filtration. Nuclei were then processed using the Chromium Single Cell ATAC v2 kit (10x Genomics) for GEM generation, barcoding, and library preparation. Oligonucleotide sequences, Tn5 loading conditions, and integrations-per-cell metrics for indexed Tn5 have been described previously [3, 49].

### Library preparation and sequencing

Multiplexed libraries were prepared as described previously [50] and sequenced at the DKFZ NGS Core Facility on the Illumina NovaSeq 6000 v1.5 platform with SP or S4 flow cells and 100 bp (RNA, ADT, ATAC) or with 250 bp paired-end reads (VDJ libraries).

### Implementation of the Split-flow pipeline

Split-flow was implemented in Nextflow DSL2 [51]. The pipeline takes as input Cell Ranger output directories and a sample sheet specifying the experimental design (sample-to-tag mapping, number of donors per channel). Each sample given in the input file is processed independently through cell calling, demultiplexing, doublet detection, and QC visualization, producing multimodal objects in both R/Seurat and Python/Scanpy formats. All tools run in Singularity containers to ensure reproducibility. For further information please refer to https://github.com/RippeLab/split-flow.

### Cell calling

Cell calling was performed using both Cell Ranger (v7.1.0) and CellBender (v0.3.0) [12, 16]. CellBender remove-background was applied to the raw count matrices to distinguish cell-containing droplets from empty droplets while correcting for ambient RNA contamination. The filtered .h5 output was converted to Seurat-readable format using Pytables and imported via Read10X_h5(). Cell Ranger and CellBender cell calls were compared by examining library size distributions for RNA and ADT counts to identify intermediate-complexity droplets recovered by CellBender but classified as empty by Cell Ranger. For the work described in this manuscript, cell calling was primarily done by CellRanger-multi for the MM dataset, and the downstream operations were applied on the union barcode set received on the independent CellRanger-multi runs of the downsampled objects, achieved using the built-in subsample_rate parameter at rates of 0.01, 0.1, and 0.5, as well as the full dataset, to enable comparability. Cell calling for the AML snMultiome dataset was performed using Cell Ranger ARC (v2.0.2). Cell Ranger Multi (v7.1.0) was run in parallel to generate CMO count matrices, with the filtered matrices subsequently restricted to Cell Ranger ARC–called barcodes to ensure a consistent cell set across modalities. Cell calling for the AML scRNA-seq dataset was performed using Cell Ranger Multi (v7.1.0). Chromatin accessibility data were processed independently using Cell Ranger ATAC (v2.1.0).

### Demultiplexing

Tag-based demultiplexing was performed per Chromium channel using HTODemux() and MultiSeqDemux() in Seurat [52]. For SNP-based demultiplexing, genotype-free modes of Souporcell [40] and cellsnp-lite followed by Vireo [39] were run using the Demuxafy Singularity image [43]. Position-sorted BAM files from Cell Ranger and union barcode lists from the downsampled CellRanger runs served as input.

Common SNPs from the 1000 Genomes Project filtered and provided by Demuxafy (GRCh38_1000G_MAF0.01_ExonFiltered_ChrEncoding.vcf) was used to pile up SNPs and during the pileup, variants were filtered further with MAF filtering using 0.05. Vireo was then executed on the resulting cellsnp-lite output with the expected donor number set to the number of pooled samples for that library and a fixed random seed.

For SNP based demultiplexing in genotype-aware mode, germline variants were called against the hg38 reference using strelka2 for 11 out of 14 total donors. Next, variants were annotated using variant effect predictor and were filtered for high quality (PASS filter). Individual .vcf files were then merged using bcftools merge with missing genotypes set to homozygous reference using the parameter –missing-to-ref. The merged file was filtered for biallelic SNPs only and an partial genotype-aware vireo run was performed setting the expected number of donors to 14 and submitting a .vcf reference with the present 11 donor alone. This step allowed inferring of the missing genotypes of the three donors anonymously. The genotypes of these three patients were extracted from GT_donors.vireo.vcf.gz, subsetted to the SNP panel derived from the original 11-donor merged VCF using bcftools view -R and merged using bcftools merge again. The final merged .vcf file containing SNPs derived from 11 patients and 3 inferred ones was used to run the final vireoWGS run.

### Doublet detection

Heterotypic doublet detection was performed using scds [28] and scDblFinder [25] on the non-negative droplets. Cells flagged as doublets by both transcription-based doublet detection algorithms were later marked in the decision making framework as txn_doublet. Cell type annotation was applied using Azimuth with human bone marrow reference (bonemarrowref) which assigns cell types with two level of resolution [53]. In this manuscript, only predicted.celltype.l1 annotations were utilized for simplicity purposes.

### Split-flow decision making logic

We first classify droplets based on concordant global assignment status across demultiplexing methods assigning each to one of the following categories: confident singlet, confident doublet, confident unassigned, potential singlet/doublet/unassigned (majority agreement), or not confident (no majority vote callable). Donor assignment is also evaluated across all demultiplexing methods by using a majority vote like strategy and quantifying the level of support for the majority vote, distinguishing between donor calls supported by multiple methods, a single SNP-based demultiplexing method, or hashing alone. Cells for which majority vote cannot be retrieved, and donor signals cannot be resolved are labeled as non-recoverable, with further distinction based on whether donor evidence exists (i.e. tie cases). In parallel, we derive a transcriptional doublet status per droplet by combining the outputs of two transcriptome-based doublet detection tools used in Split-flow, (scds and scDblFinder respectively) labeling each as a transcriptomic singlet, doublet, or not confident when the two methods disagree.

To derive final assignments, we combine this information into an intermediate call that captures agreement or conflict between modalities. Droplets are then assigned to five final categories using a rule-based prioritization framework (Figure 4), implemented as a rule-based (case_when in R) scheme rather than a pure decision tree. Transcriptional doublet evidence has highest priority. Remaining droplets are resolved based on donor assignment strength and demultiplexing consensus, yielding high-confidence singlets, low-confidence singlets, doublets, unassigned, and donor-assignable but non-recoverable cells. The latter captures droplets with inconsistent donor evidence that should be evaluated with orthogonal data.

### TCR sequence data analysis and validation

TCR sequences were processed using the scRepertoire framework and custom functions for strategy-based concordance metrics. Clonotypes were defined based on the strict TRB CDR3 amino acid sequence (CTaa). For each demultiplexing strategy, three complementary metrics were derived. First, T-cell yield was defined as the number of cells with a valid TCR annotation retained after filtering. Second, clonotype purity was quantified at the clonotype level as the fraction of cells within a clonotype assigned to the dominant donor; clonotypes detected across multiple donors were considered impure, and the percentage of such clonotypes was reported per strategy. Third, to assess biological plausibility of cross-donor sharing, shared clonotypes (those appearing in more than 2 donors within a pool) were compared against curated human VDJdb CDR3β sequences (downloaded April 2026)[37, 38]. Precision was defined as the proportion of shared clonotypes with a VDJdb match; absolute counts of confirmed versus total shared clonotypes were reported per strategy.

### Integration of datasets and multiomics readouts

Seurat objects from 8 channels (same pool loaded independently to 8 10x chip channel) were loaded and merged into a single Seurat object using merge function via defining channel number as sample identity in the MM dataset. The merged object was filtered for cells called only in the full dataset, RNA and ADT modalities were processed independently. Gene expression counts derived from CellRanger-multi were normalized using SCTransform with default parameters and 3000 highly variable gene selection, followed by a PCA applied on the SCT normalized assay. ADT counts were normalized using centered log-ratio (CLR) normalization (margin set to 2), and were subjected to PCA using all available features (n=277). Elbow plots on 50 PCs informed the selection of the top 30 PCs for the RNA and 25 PCs for the ADTs. A weighted nearest neighbor (WNN) analysis was performed using FindMultiModalNeighbors() in Seurat using 30 and 25 dimensions respectively for RNA and ADT in order to integrate RNA and ADT modalities [52]. A joint WNN UMAP embedding was computed from the WNN graph and leiden clustering was applied on the WSNN graph using three different resolutions 0.4, 0.5, and 0.6. UMAP visualizations were split to each channel. For comparing the impact of singlet definition strategy, three separate objects were created by filtering the main merged object into (1) singlets identified by hash demultiplexing alone, (2) concordant singlets identified by all demultiplexing methods (vireo, vireoWGS, hash, souporcell) and (3) union singlets identified by at least one of the demultiplexing strategies as a singlet. SCT normalization on CellRanger gene expression counts, PCA, nearest neighbour graph construction using the top 30 PCs were applied independently on each singlet set and a UMAP embedding was computed using top 30 PCs.

Seurat objects from two pools of the AML snMultiome dataset were merged and the chromatin accessibility assay was added using Signac with EnsDb.Hsapiens.v86 annotation. RNA and ATAC modalities were processed independently, with RNA log-normalised using 3,000 highly variable features and 50 PCs, and ATAC TF-IDF normalised with truncated SVD (50 components, LSI component 1 excluded), both batch-corrected using Harmony. Weighted nearest neighbor (WNN) integration was performed using the top 50 RNA and components 2-50 of the ATAC reduction, and cell type annotation was performed using Azimuth with the bone marrow reference.

For the AML scRNA-seq and snATAC-seq datasets, RNA and ATAC modalities were processed independently. Seurat objects from three pools were merged, log-normalized with 3,000 highly variable features, scaled with mitochondrial percentage regressed out, and batch-corrected using Harmony on the top 50 PCs. Per-sample Signac objects from Cell Ranger ATAC output were merged, TF-IDF normalized, and dimensionality reduction was performed using truncated SVD on LSI components 2-20.

### Software

Software versions used in this study are listed in **Supplementary Table 1**.

## Author contributions

Conceptualization: ES, SSt, KR. Methodology: ES, SSt, MB. Software: ES, SSt, MB. Investigation: ES, SSt, MB, KR. Resources: NP, SSc, MF, NW, ML, HD, KD, MR, JPM, KR. Writing – original draft: ES, KR. Writing – review & editing: ES, SSt, MB, KR. Visualization: ES, SSt, MB. Supervision: SS, OS, KR.

## Acknowledgements

We thank Florian Heyl and Marc Zapatka for help and discussion, and the Sample Processing Lab (SPL), the High Throughput Sequencing unit of the Genomics & Proteomics Core Facility and the Omics IT, Data Management Core Facility (ODCF) of the German Cancer Research Center (DKFZ), the Biobank Multiple Myeloma UKHD and the Myeloma Registry for excellent services. This work was supported by German Research Foundation (DFG) grants SFB1074 (B12, Z01), RI 1283/15-2 (K.R.), LU 429/16-2 (M.L.) and MA 7792/1-2 (J.-P.M.) within Research Group FOR2674 and TRR179 (Z03), and by the German Federal Ministry of Education and Research (BMFTR) via project SATURN3 (01KD2206B) within the National Decade against Cancer program. N.P. was supported by a European Union Horizon Europe Marie Skłodowska-Curie Postdoctoral Fellowship (101068158) and the Olympia Morata Program of Heidelberg University.

## Declarations

### Ethics approval and consent to participate

This study was conducted following the approval of the ethics committees at the Medical Faculty of Heidelberg University, University Hospital Freiburg, and University Hospital Ulm. All experiments in this study involving human tissue or data adhered to the Declaration of Helsinki and relevant national and international ethical guidelines.

### Consent for publication

Not applicable.

### Competing interests

The authors declare that they have no competing interests.

### Availability of data and analysis scripts

Split-flow is provided via a Github repository at https://github.com/RippeLab/Split-flow. The software used for the data analysis is listed in **Supplementary Table 1**.

## Abbreviations

ADT: Antibody-derived tag
CITE-seq: cellular indexing of transcriptomes and epitopes by sequencing
HTO: hashtag oligonucleotide
scRNA-seq: single-cell RNA sequencing
SNP: single nucleotide polymorphism
TCR: T cell receptor
TCR-seq: T cell receptor sequencing
UMAP: uniform manifold approximation and projection
UMI: unique molecular identifier

## Notes

### Competing Interest Statement

The authors have declared no competing interest.

