## Supplementary material for "Disagreement between demultiplexing methods reveals structured cell quality gradients in multiplexed single-cell data"

#### Content

**Additional files: Fig. S1-S6**

Fig. S1. Effect of sequencing depth on demultiplexing performance.
Fig. S2. HTO signal quality and doublet detection concordance.
Fig. S3. Intermediate assignment categories map to final decision classes.
Fig. S4. Split-flow workflow in AML Multiome data.
Fig. S5. Split-flow workflow in AML scRNA-seq and scATAC-seq data.
Fig. S6. Tn5-based demultiplexing and quality assessment of AML scATAC-seq data.

**Additional files: Table S1**

Table S1. Software utilized with corresponding versions and sources.

### Additional files: Fig. S1-S6

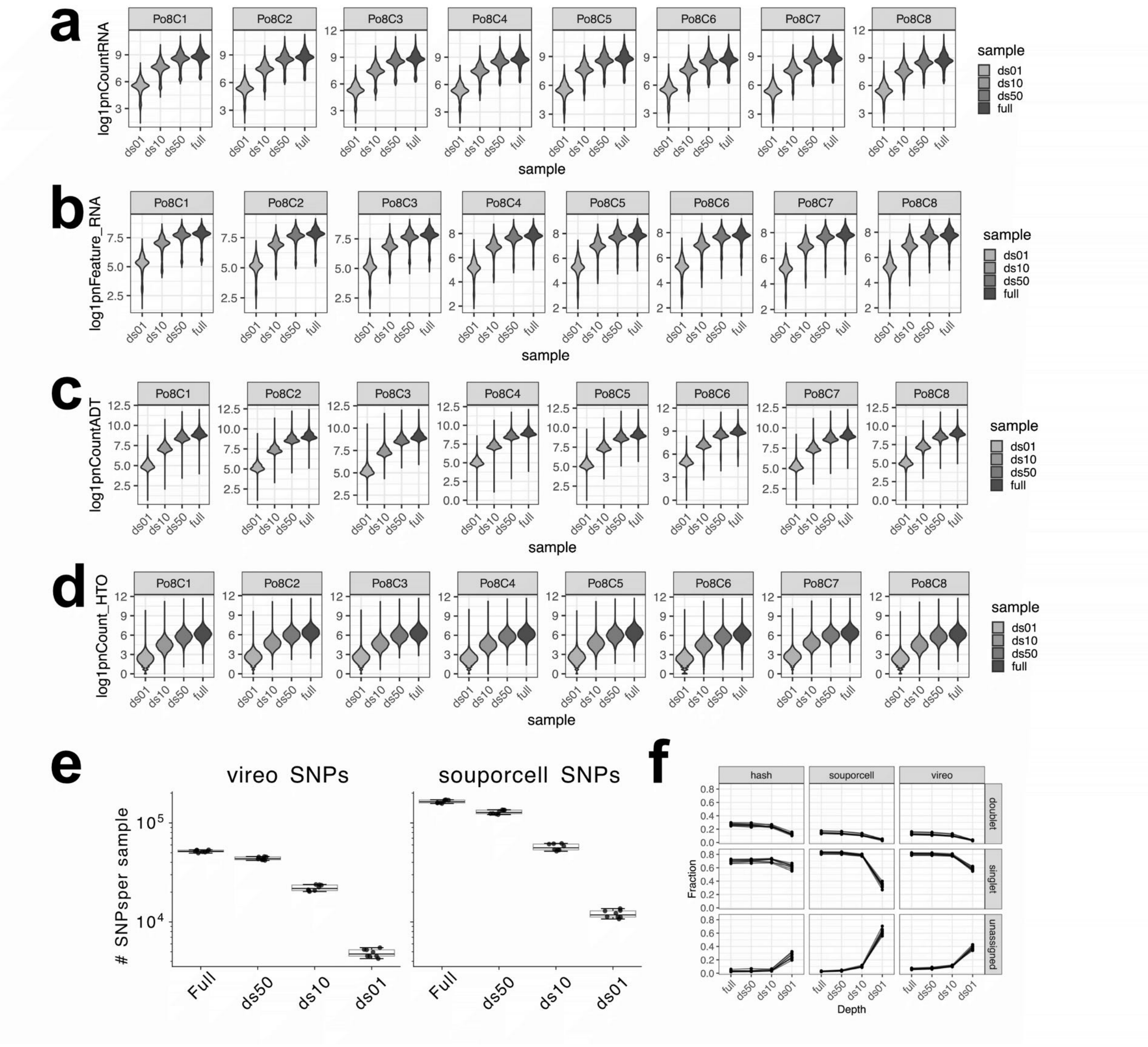


**Supplementary Figure S1. Effect of sequencing depth on demultiplexing performance.**
(**a**) Distribution of per-cell QC metrics across eight Chromium channels (PolBC1–PolBC8) at full sequencing depth and after downsampling to 50%, 10%, and 1% of total reads (ds50, ds10, ds01). Violin plots with overlaid dot plots show the per-cell distribution within each channel. Metrics shown are: log1p-transformed total RNA counts (nCount_RNA). **(b)** log1p-transformed number of detected RNA features (nFeature_RNA). **(c)** log1p-transformed total ADT counts (nCount_ADT). **(d)** log1p-transformed total HTO counts (nCount_HTO). **(e)** Number of informative SNPs detected at full depth and after downsampling (ds50, ds10, ds01) for the cellsnp-lite + Vireo and Souporcell pipelines. **(f)** Fraction of cells classified as singlet, doublet, or unassigned across sequencing depths for HTODemux, Souporcell, and Vireo. Lower sequencing depth increases the fraction of unassigned cells and alters singlet/doublet proportions, with method-specific sensitivity.

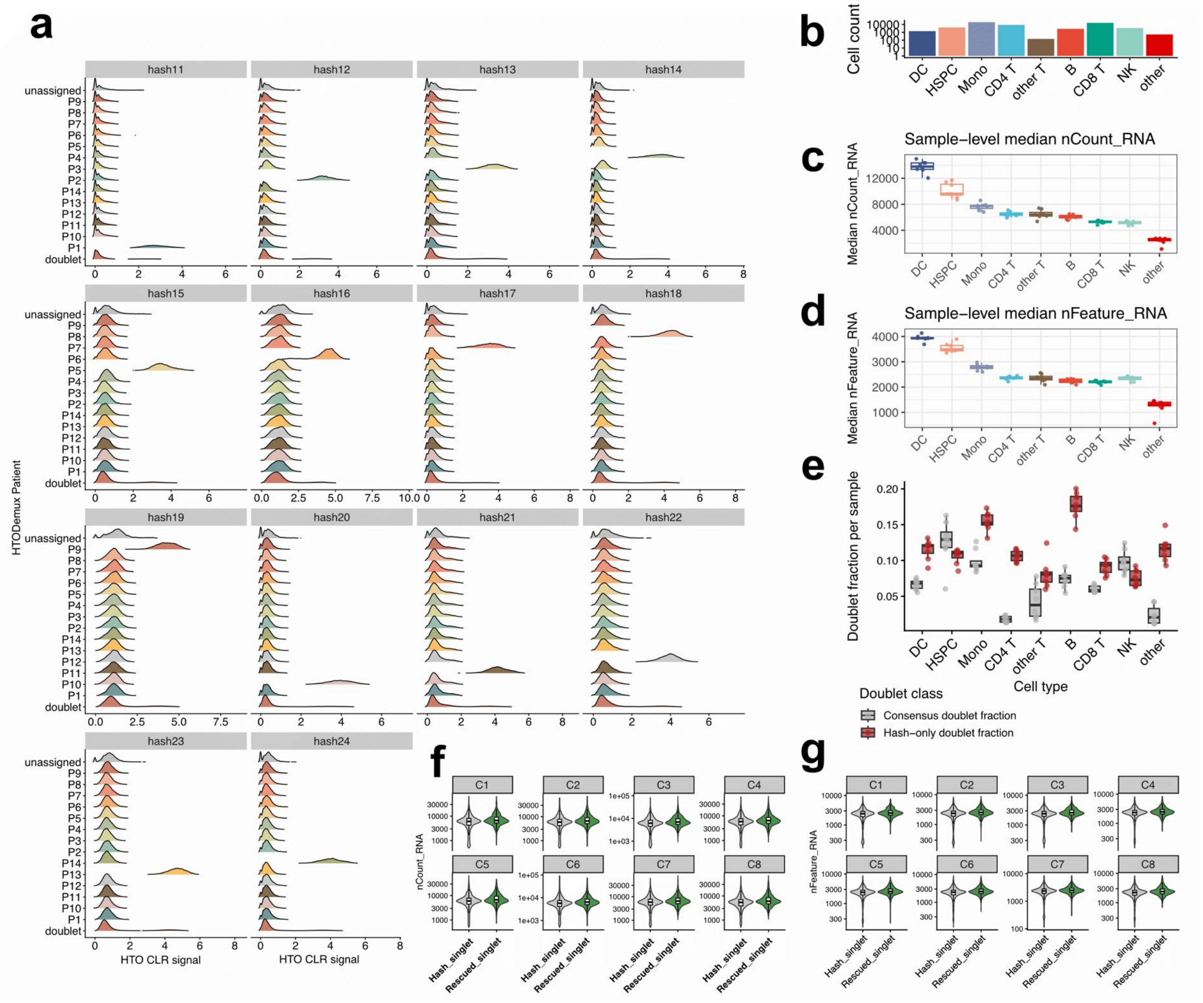


**Supplementary Figure S2. HTO signal quality and doublet detection concordance.**

(**a**) Ridge density plots of CLR-normalized HTO signal for each hashing antibody (Hash-11 to Hash-24), stratified by HTODemux-assigned patient identity. Each ridge represents the per-cell HTO signal distribution within a donor group; facets correspond to individual hash features. Clear separation between signal-positive and signal-negative populations indicates efficient sample barcoding. (**b**) Bar plot showing the number of cells per BM Azimuth level-1 cell type in the consensus singlet barcode set. The y-axis is log-scaled. (**c**) Boxplots showing the distribution of median total RNA counts across BM Azimuth cell types in the consensus singlet set; points represent individual channels. (**d**) Boxplots showing the distribution of median total detected gene counts across BM Azimuth cell types in the consensus singlet set; points represent individual channels. (**e**) Boxplots of per-sample doublet fractions stratified by cell type, comparing consensus doublets (cells classified as doublets by all five methods: HTO, Souporcell, Vireo, scds, and scDblFinder) and hash-only doublets (cells classified as doublets exclusively by HTO-based demultiplexing). Points represent individual channels. (**f**) Comparison of total RNA count distributions across cells in consensus singlets classified as singlets by all five methods (HTO, Souporcell, Vireo, scds, and scDblFinder) and SNP-rescuable hash doublets. (**g**) Comparison of detected gene count distributions across cells in consensus singlets classified as singlets by all five methods (HTO, Souporcell, Vireo, scds, and scDblFinder) and SNP-rescuable hash doublets.

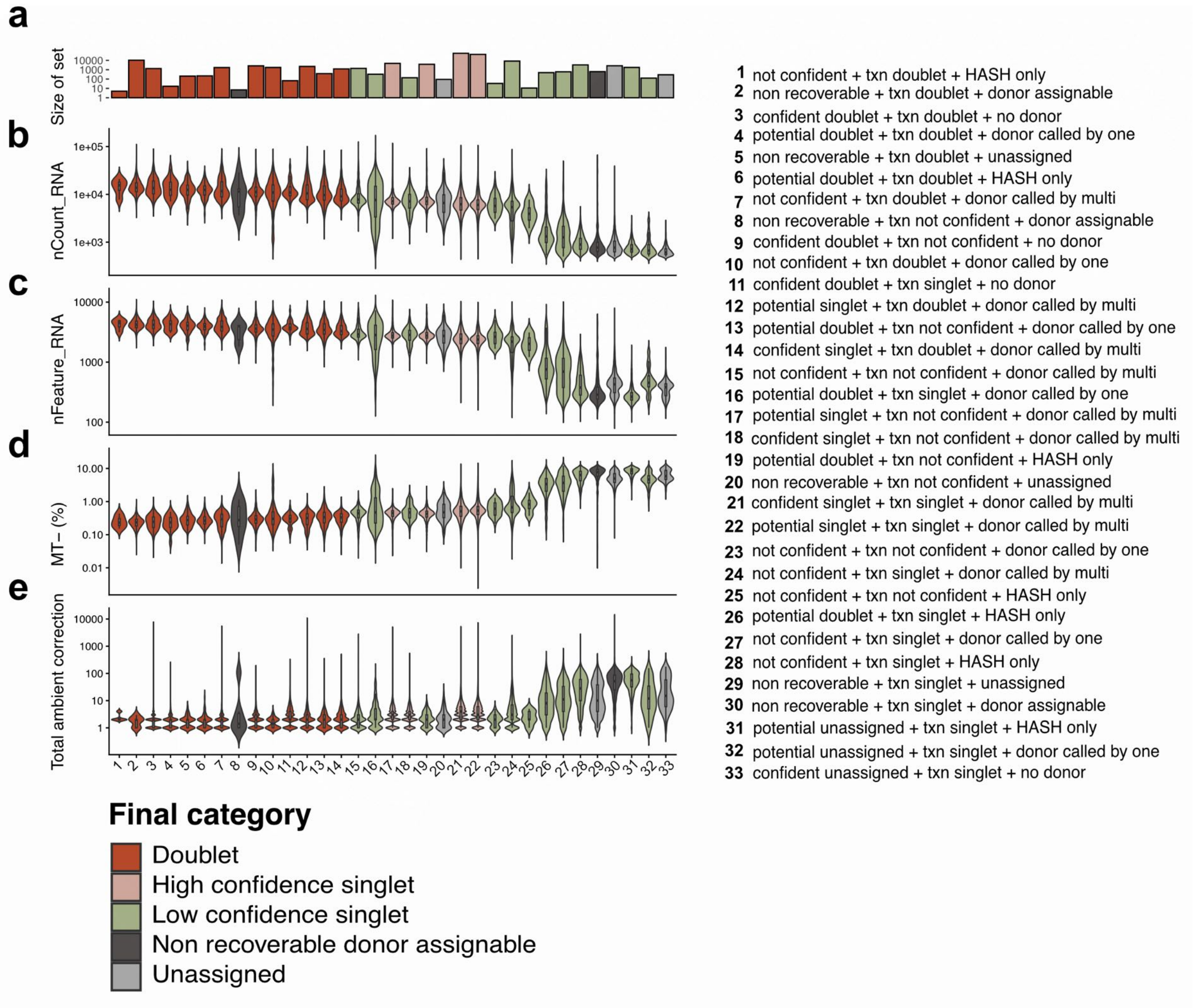


**Supplementary Figure S3. Intermediate assignment categories map to final decision classes**. (**a**) Set size of each category in cell numbers. **(b)** Violin plots alongside intermediate categories showing total UMI counts. Boxplots are overlaid on the violins, and colors represent the final category mapping of each intermediate class. Violin plots showing the distribution of unique gene counts per cell are shown in **(c),** mitochondrial percentage per cell in **(d),** and total ambient RNA correction per cell by CellBender in **(e)**. The legend on the right explains the numbered intermediate classes. Each class is defined by three decisions. The first decision represents the majority vote on the assignment status of a droplet: confident singlet, confident unassigned, confident doublet, potential singlet, potential doublet, or potential unassigned. When there is a tie, the droplet is assigned "not confident". The second decision comes from transcriptional doublet detection tools. If both methods classify the droplet as a doublet or singlet, it is assigned "txn_doublet"/"txn_singlet"; if there is no consensus, the droplet is assigned "txn not confident". The third decision concerns donor assignment. If a donor can be assigned by majority vote, the droplet is labeled "donor called by multi". If only hashing methods assigned a donor, the droplet is labeled "HASH only". If only a single non-hashing method assigned a donor, the droplet is labeled "donor called by one". If neither majority voting nor a single method can resolve the donor (tie cases or conflicting assignments), the droplet is assigned to the category "non recoverable, txn …, donor assignable".

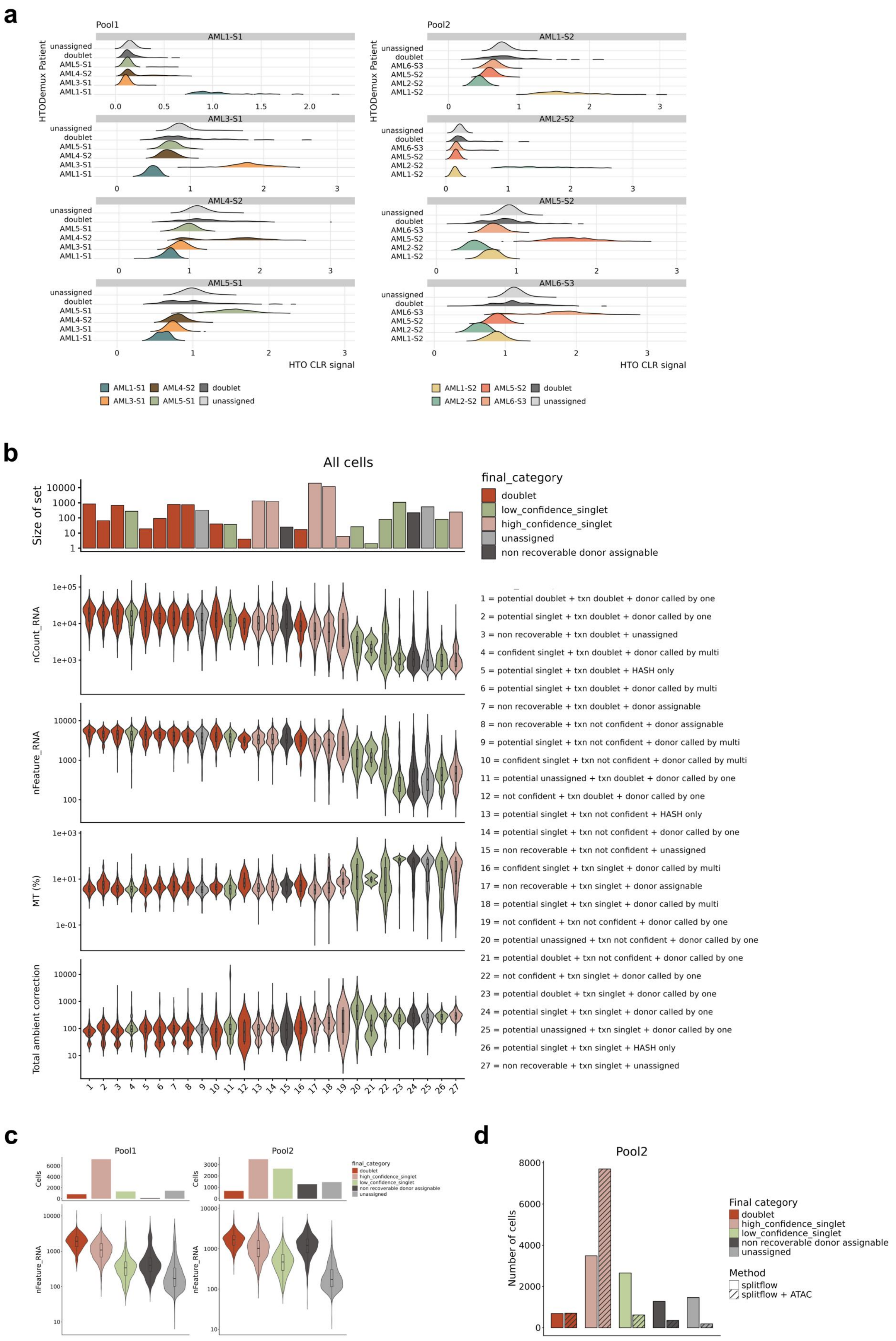
a
Pool1
Pool2
AML1-S1
AML3-S1
AML4-S2
AML5-S1
AML1-S2
AML2-S2
AML5-S2
AML6-S3
unassigned
doublet
HTODemux Patient
HTO CLR signal
AML1-S1 AML4-S2 doublet
AML3-S1 AML5-S1 unassigned
AML1-S2 AML5-S2 doublet
AML2-S2 AML6-S3 unassigned
b
All cells
Size of set
final_category
doublet
low_confidence_singlet
high_confidence_singlet
unassigned
non recoverable donor assignable
nCount_RNA
nFeature_RNA
MT (%)
Total ambient correction
1 = potential doublet + txn doublet + donor called by one
2 = potential singlet + txn doublet + donor called by one
3 = non recoverable + txn doublet + unassigned
4 = confident singlet + txn doublet + donor called by multi
5 = potential singlet + txn doublet + HASH only
6 = potential singlet + txn doublet + donor called by multi
7 = non recoverable + txn doublet + donor assignable
8 = non recoverable + txn not confident + donor assignable
9 = potential singlet + txn not confident + donor called by multi
10 = confident singlet + txn not confident + donor called by multi
11 = potential unassigned + txn doublet + donor called by one
12 = not confident + txn doublet + donor called by one
13 = potential singlet + txn not confident + HASH only
14 = potential singlet + txn not confident + donor called by one
15 = non recoverable + txn not confident + unassigned
16 = confident singlet + txn singlet + donor called by multi
17 = non recoverable + txn singlet + donor assignable
18 = potential singlet + txn singlet + donor called by multi
19 = not confident + txn not confident + donor called by one
20 = potential unassigned + txn not confident + donor called by one
21 = potential doublet + txn not confident + donor called by one
22 = not confident + txn singlet + donor called by one
23 = potential doublet + txn singlet + donor called by one
24 = potential singlet + txn singlet + donor called by one
25 = potential unassigned + txn singlet + donor called by one
26 = potential singlet + txn singlet + HASH only
27 = non recoverable + txn singlet + unassigned
c
Pool1
Pool2
Cells
nFeature_RNA
final_category
doublet
high_confidence_singlet
low_confidence_singlet
non recoverable donor assignable
unassigned
d
Pool2
Number of cells
Final category
doublet
high_confidence_singlet
low_confidence_singlet
non recoverable donor assignable
unassigned
Method
splitflow
splitflow + ATAC

e

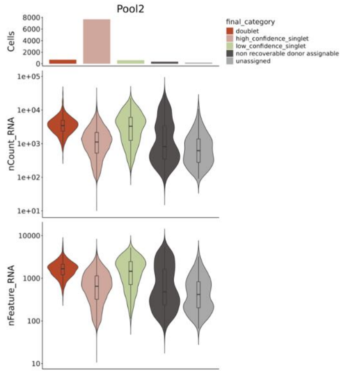


f

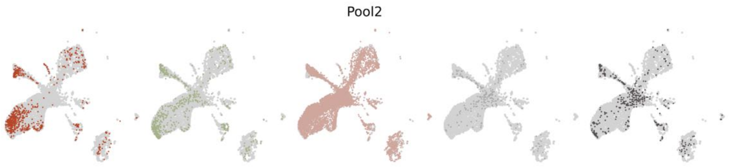


**Supplementary Figure S4. Split-flow workflow in AML Multiome data.**
(**a**) Ridge density plots of CLR-normalized HTO signal for each hashing antibody, stratified by HTODemux-assigned donor identity. Each ridge represents the per-cell HTO signal distribution within a donor group, and facets correspond to individual hashing antibodies. (**b**) Intermediate assignment categories mapped to final decision classes. A bar plot showing the number of cells in each intermediate category (1–27). Violin plots show per-cell distributions of total UMI counts (nCount_RNA), unique gene counts (nFeature_RNA), mitochondrial percentage, and ambient RNA correction estimated by CellBender across intermediate categories, with boxplots overlaid. Colors indicate the final category assigned to each intermediate class. (**c**) Bar plots showing the total number of cells assigned to each final classification category for each pool based on the Split-flow decision framework (Fig. 4) (top), and violin plots showing per-cell distributions of unique gene counts (nFeature_RNA) across these categories for each pool (bottom). (**d**) Bar plots showing the number of cells in each final classification category for Pool2 across two Split-flow configurations: one including Souporcell (RNA) and one including Souporcell (ATAC) for demultiplexing. (**e**) Bar plots showing the total number of cells assigned to each final classification category for Pool2 using Split-flow with Souporcell (ATAC) for demultiplexing (top), and violin plots showing per-cell distributions of total UMI counts (nCount_RNA) (middle) and unique gene counts (nFeature_RNA) (bottom) across these categories. (**f**) UMAP embedding colored by final classification category for Pool2 using Split-flow with Souporcell (ATAC) for demultiplexing.

a

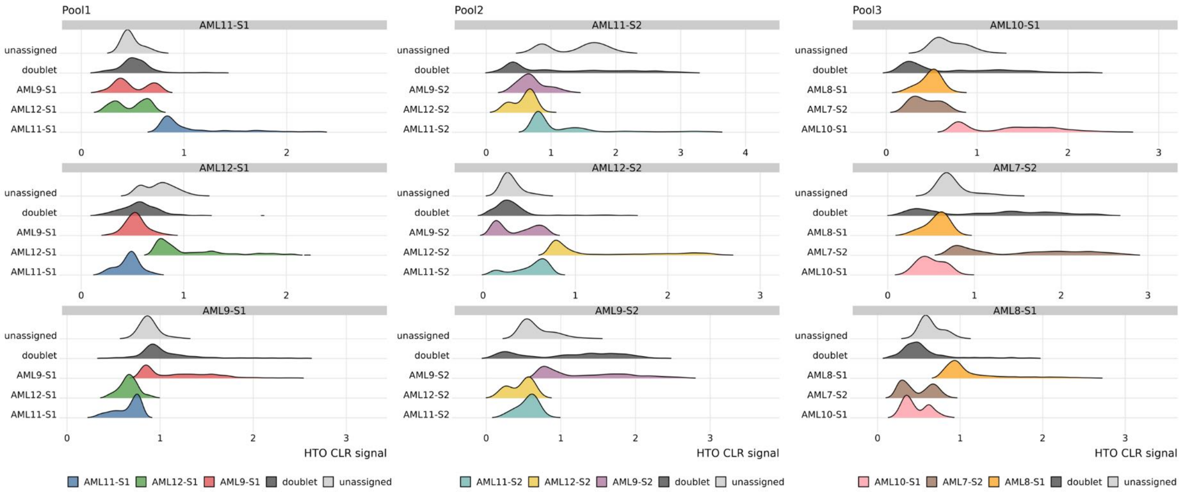


b

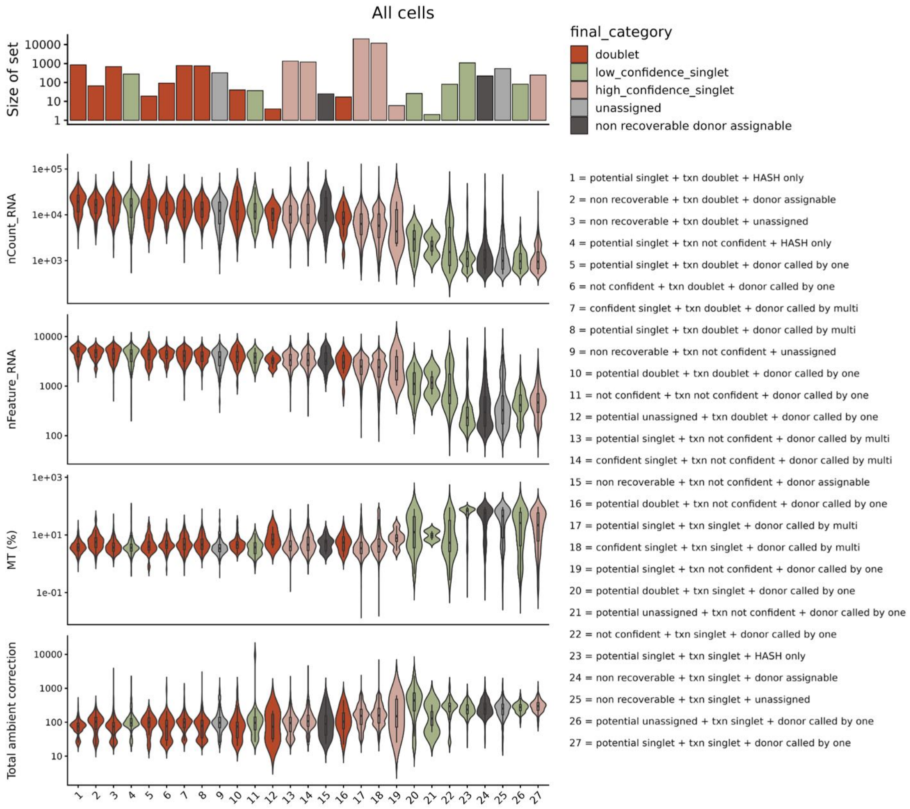


c

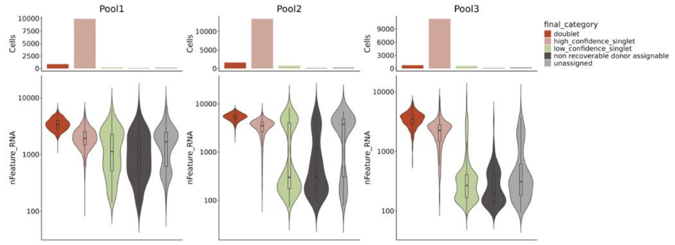


**Supplementary Figure S5. Split-flow workflow in AML scRNA-seq and scATAC-seq data.** **(a)** Ridge plots showing CLR-normalized HTO signal distributions for each hashing antibody, grouped by HTODemux-assigned donor identity. Each ridge represents the distribution of HTO signal across cells within a donor group, with separate facets corresponding to individual hashing antibodies. **(b)** Mapping of intermediate assignment categories to final decision classes. Bar plot indicating the number of cells in each intermediate category (1–27). Violin plots display per-cell distributions of total UMI counts (nCount_RNA), detected gene counts (nFeature_RNA), mitochondrial percentage, and ambient RNA correction estimated by CellBender across intermediate categories, with boxplots overlaid. Colors denote the final class assigned to each intermediate category. **(c)** Bar plots summarizing the number of cells assigned to each final classification category per pool according to the Split-flow decision framework (Fig. 4) (top), alongside violin plots showing per-cell distributions of detected gene counts (nFeature_RNA) across these categories (bottom).

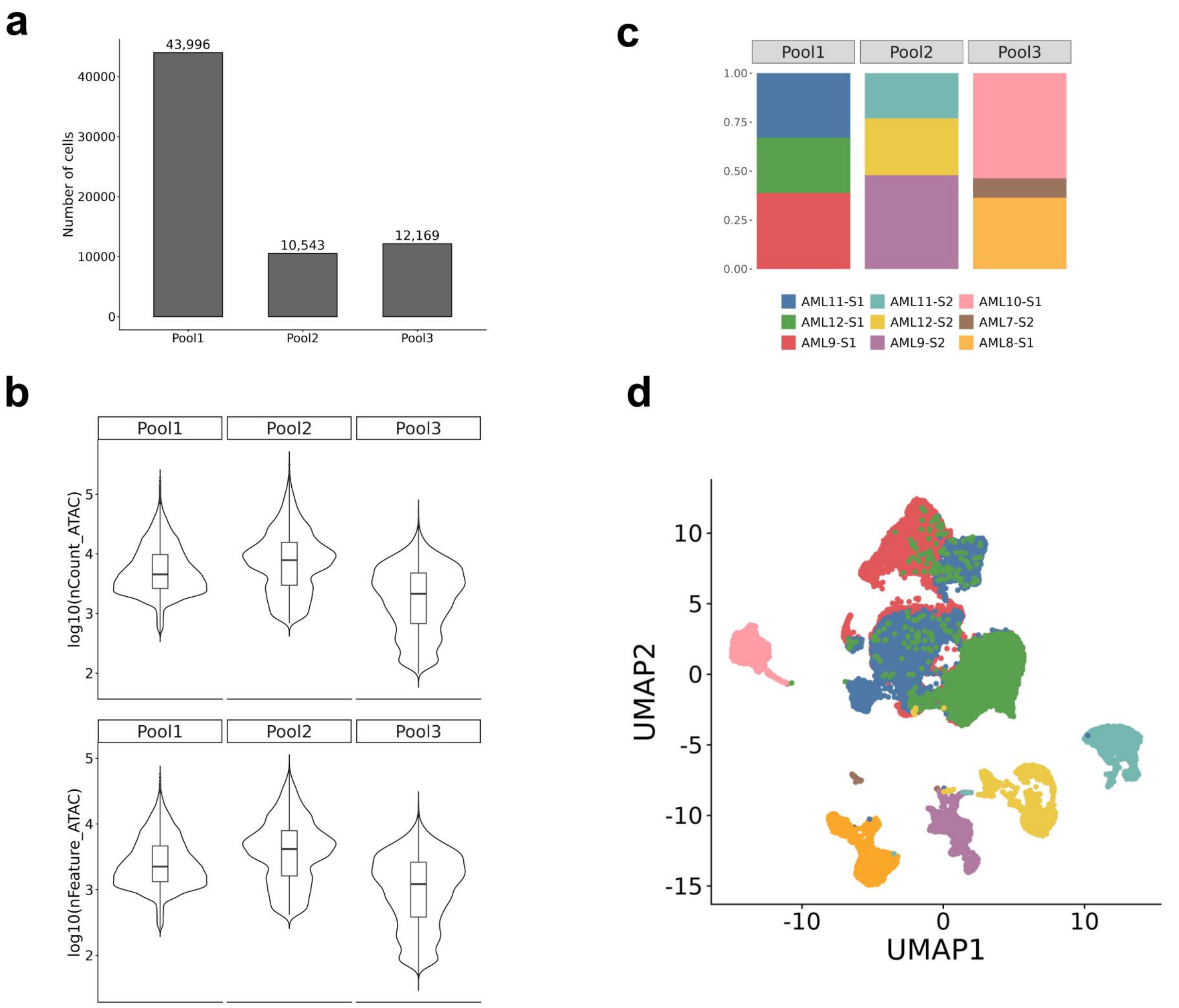


**Supplementary Figure S6. Tn5-based demultiplexing and quality assessment of AML scATAC-seq data.** (**a**) Bar plot showing the total number of cells detected by Cell Ranger in each pool. (**b**) Violin plots summarizing quality control metrics across pools, including total ATAC fragment counts (nCount_ATAC) and the number of accessible features (nFeature_ATAC). (**c**) Stacked bar plots showing the proportion of cells assigned to each donor within each pool based on Tn5 barcoding. (**d**) UMAP embedding of cells colored by donor identity. Only cells with nCount_ATAC between 3,000 and 100,000 were included.

### Supplementary Tables

**Supplementary Table 1. Software utilized with corresponding versions and sources**

| Software | Version | Source |
|---|---|---|
| **CellBender** | v.0.3.0 | [1] |
| **Cell Ranger** | v.7.1.0 | [2] |
| **cellSNP-lite** | v1.2.4 | [3] |
| **Demuxafy** | v.2.0.1 | [4] |
| **dplyr** | v.1.1.4 | [5] |
| **dsb** | v.1.0.3 | [6] |
| **Muon** | v.0.1.5 | [7] |
| **Nextflow** | v.23.10.0 & v.22.10.7 | [8] |
| **Pytables** | v.3.8.0 | [9] |
| **Python** | v. 3.9.18 & v3.1.6 | https://www.python.org |
| **R** | v.4.3.2 | https://www.r-project.org |
| **qs (R)** | v.0.25.7 | [10] |
| **Scanpy (Python)** | v.1.9.8 | [11] |
| **scDblFinder (R)** | v.3.10 | [12] |
| **Scds (R)** | v.1.17.0 | [13] |
| **scTransform (R)** | v.0.4.1 | [14] |
| **scRepertoire** | v.2.6.2 | [15] |
| **scvi-tools** | v.1.0.3 | [16, 17] |
| **Seurat (R)** | v.5.0.1 | [18] |
| **Souporcell** | v2.0 | [19] |
| **tidyr (R)** | v.1.3.1 | [20] |
| **Vireo** | v0.5.8 | [21] |
